# Culture-dependent identification of rare marine sediment bacteria from the Gulf of Mexico and Antarctica

**DOI:** 10.1101/2024.06.11.598530

**Authors:** Sarah J. Kennedy, Celine Grace F. Atkinson, Tristan J. Tubbs, Bill J. Baker, Lindsey N. Shaw

**Author notes:** **Correspondence:** Sarah J. Kennedy, Lindsey N. Shaw.

## Abstract

Laboratory-viable cultivars of previously uncultured bacteria further taxonomic understanding. Despite many years of modern microbiological investigations, the vast majority of bacterial taxonomy remains uncharacterized. While many attempts have been made to decrease this knowledge gap, culture-based approaches parse away at the unknown and are critical for improvement of both culturing techniques and computational prediction efficacy. To this end of providing culture-based approaches, we present a multi-faceted approach to recovering marine environmental bacteria. We employ combinations of nutritional availability, inoculation techniques, and incubation parameters in our recovery of marine sediment-associated bacteria from the Gulf of Mexico and Antarctica. The recovered biodiversity spans several taxa, with 16S-ITS-23S rRNA gene-based identification of multiple isolates belonging to rarer genera increasingly undergoing phylogenetic rearrangements. Our modifications to traditional culturing techniques have not only recovered rarer taxa, but also resulted in the recovery of biotechnologically promising bacteria. Together, we propose our stepwise combinations of recovery parameters as a viable approach to decreasing the bacterial knowledge gap.

## Introduction

Bacteria, the most abundant form of life on Earth, continue to evade taxonomic understanding. Since the advent of modern microbiology, we have yet to categorize and study more than 1% of the total predicted biodiversity (1). The discovery and study of this cryptic bacterial majority would undoubtedly provide critical advancements within many fields including (but not limited to) pharmaceutical, biotechnological, and agricultural (2, 3). While there are hypotheses as to why these bacteria remain outside culture-capabilities, our focus remains on how to elicit their growth within the laboratory.

Environmental bacteria must overcome many obstacles before reaching viability within the laboratory. The transition from environment to laboratory often requires alteration to typical bacterial culturing campaigns that commonly prioritize quantity over taxonomic quality. Many successful culture-based approaches bridge the gap via provision of generalized/high-throughput growth requirements balanced with physicochemically-individualized recovery programs. These have included changing media gelling agents (4, 5), decreasing incubation temperatures and nutrient availability (6), alongside myriad combinations of proven, traditional culturing techniques (7). Indeed, conventional fecal sample preparation, inoculation, and incubation on commercially available growth media has resulted in recovery of novel human gut bacteria, as benchmarked by 16S rRNA gene analysis (8).

While traditional recovery methods have existed since the advent of modern microbiology, there are undeniable advantages to thinking outside the Petri plate. One such innovation, the iChip, disregards the Petri plate and returns environmental bacteria to the sampling site, allowing unfettered, natural growth (9). This controlled return to natural conditions restores the complex communities’ diffusion-mediated metabolic/ physiological networks. The iChip’s enclosure of environmental bacteria within semi-permeable membranes allows diffusion of integral environmental/community growth factors (10) that would otherwise be disrupted upon traditional monoculture. Indeed, when examined for recovery efficiency, diffusion-enabled growth recoveries approached 50% of tested soil populations, as compared to the approximately 1% obtained via Petri plate growth (11).

While non-traditional techniques can yield novel chemotaxonomy, they are also infrequently developed and require extensive validation. Non-traditional techniques may also be more difficult to scale-up and generalize for larger sample sizes and disparate environmental conditions. It is because of these reasons, and the allure of novel chemotaxonomy, that environmental bacterial recovery is improved through the application of many different combinations of approaches. Traditional selective recovery methods still have a chance of avoiding bacterial rediscoveries, with some holistic approaches succeeding in recovering highly biodiverse taxa (12). Furthermore, examination of geographically unique or otherwise under-examined environments has been shown to decrease taxonomic rediscovery. One of the greatest under-examined resources is the marine environment. Although marine sediments encompass ∼70% of the planet’s surface (13), more than 80% of the ocean remains unexplored (14). Within those underexplored niches, it is estimated that more than 75% of sediment genera remain uncultured (15). While culture-independent metagenomics sheds light on such potential biodiversity, culture-dependent approaches provide functionality to bioinformatic findings (16). Indeed, culture-dependent techniques continually recover rare marine bacterial chemotaxonomy (17), providing physiological and metabolic insights for improvement of future taxonomic discoveries (18).

To the end of combining proven techniques to culture previously uncultured organisms, we present a high-throughput approach to culturing marine bacteria. Our methodology combines selective and non-selective/permissive inoculation and incubation parameters. We examine the effects that nutritional availability, inoculation technique, and incubation temperature have on laboratory recovery. Through the stepwise combinations of traditional techniques, we were able to recover 286 sediment-associated bacterial isolates from the Gulf of Mexico and Antarctica. Within the recovered taxonomy, we observe several isolates belonging to new and developing genera, indicating success towards our goal of decreasing the bacterial taxonomic knowledge gap. Collectively, our method can be adapted and expanded to other environmental niches to increase the total cultured biodiversity.

## Results and Discussion

### The application of multiple recovery conditions results in isolation of 121 Gulf of Mexico bacterial isolates from 3 phyla

We employed a balanced approach of selective factors and permissive countermeasures (see methods) to characterize the culturable bacterial biodiversity from Gulf of Mexico sediment. While seemingly contradictory, our holistic approach allowed for the successful recovery of over 120 bacterial isolates. While the permissive culturing approaches (increased incubation temperature and nutrient availability) and non-selective inoculation (sediment-stamping) were intended to minimize selection pressures during transition to viable monoculture, our stepwise selective factors (decreased incubation temperature and available nutrients, heat-treatment inoculation) were included to discourage overgrowth of quickly growing isolates collectively termed “microbial weeds.”

Through combinations of both selective and permissive inoculation, nutrient availability, and incubation temperature, we recovered a diverse subset of bacterial taxonomy from our Gulf of Mexico samples. The 16S-ITS-23S rRNA gene sequencing of recovered sediment-associated bacteria reveals that isolates span three taxonomic phyla and twenty-five genera. Within those three phyla recovered, four taxonomic classes and nine taxonomic orders were represented (**Supplemental Table S1**).

The 16S-ITS-23S rRNA BLASTn results for the Gulf of Mexico sediment-associated isolates are presented in **Supplemental Table S2.** To visually summarize the recovered phylogeny and their associated taxonomic rarity and recovery conditions, we present a phylogenetic tree of the 121 Gulf of Mexico sediment-associated isolates (**Figure 1**). In addition to the dendrogram (**Figure 1A**), we summarize the isolates’ BLASTn percent identity values to the most similar NCBI-available match (**Figure 1B**); recovery nutrient conditions (**Figure 1C**); inoculation methodologies (**Figure 1D**); and initial recovery incubation temperatures (**Figure 1E**).

**Figure 1.**
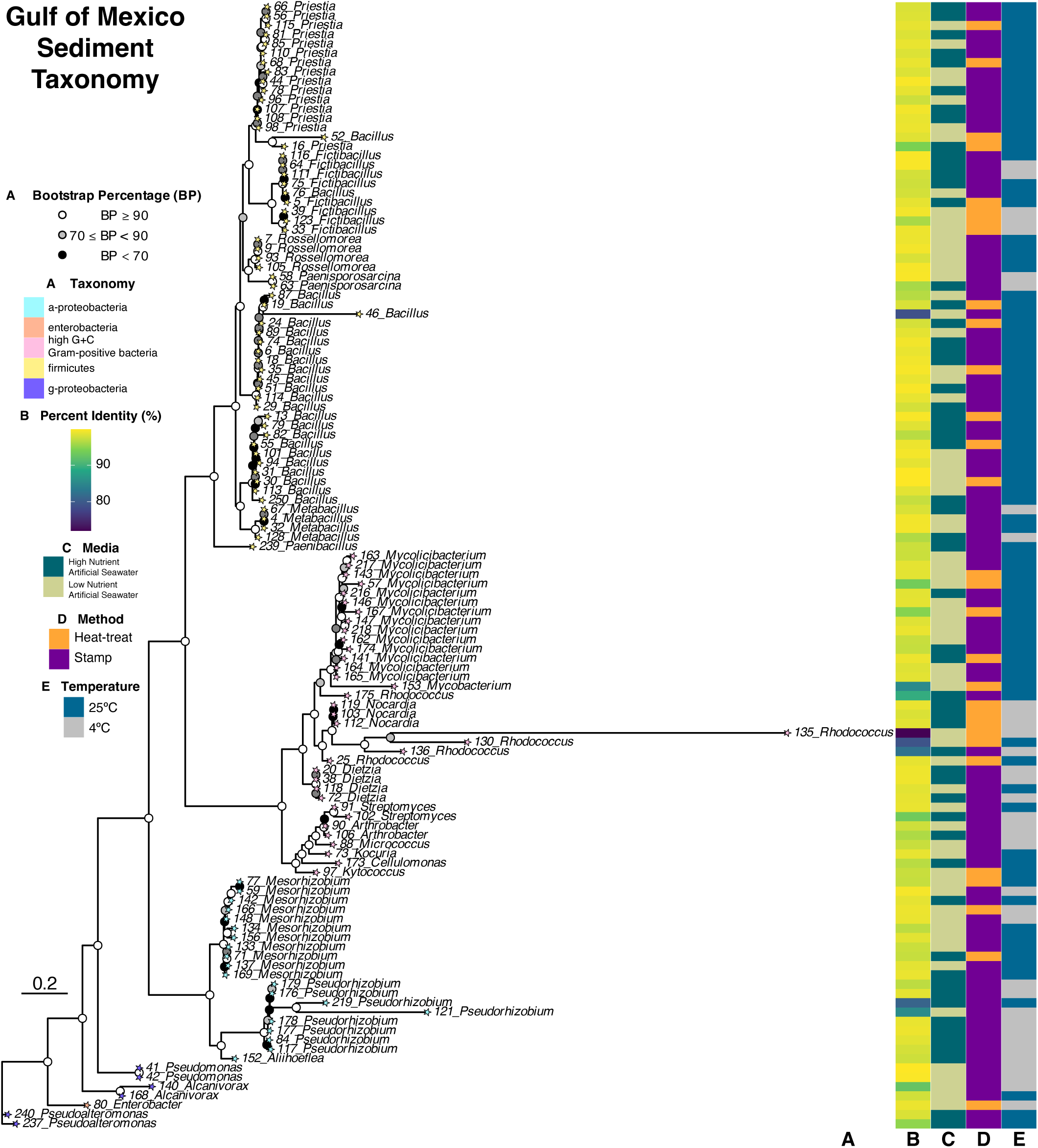
Phylogeny of recovered Gulf of Mexico sediment-associated bacteria shows varied taxa represented. The 16S-23S-rRNA sequences from 121 Gulf of Mexico isolates were compared to the 16S_ribosomal_RNA BLASTn Database for most similar genera identifications, and then aligned and plotted within a phylogenetic tree (**A**). Data is presented as in-laid maps, from left to right: BLASTn percent identity to the most similar result (**B**), recovery media nutrient availability (**C**), recovery inoculation method (**D**), and recovery incubation temperature (**E**).

As a whole, the majority of recovered isolates are Gram-positive (**Figure 1A**). Although more firmicutes (low-GC) were recovered (59 isolates; 7 genera), more Actinobacteria (high-GC) genera were isolated (35 isolates; 11 genera). The high-GC/Actinobacteria branches contain several distinct, bootstrap-supported subclusters, particularly within the taxonomic range observed in the *Mycolicibacterium* and *Rhodococcus* genera. Conversely, although nearly half of the Gulf of Mexico library were low-GC/firmicutes (59/121 isolates, ∼48.76%) the strains are closely clustered. Upon closer inspection, however, the isolates have taxonomic origins as *Bacillus* or previously-considered *Bacillus* genera (**Figure 1A**). Critically, however, the clade contains multiple smaller clusters of divergent, *Bacillus*-like taxa that could one day be re-classified.

Overall, the BLASTn-determined percent identity values (% ID) range from 71.99 – 99.89% (**Figure 1B**). As the project aimed to recover rare taxonomy, the heatmap contains 26 isolates (within 17 genera) that have % IDs <97%, indicating potential species differentiation from the closest BLASTn match (19). Notably, within those taxonomically-promising isolates, 16 isolates (within 10 genera) present % IDs <95%, indicating potential genera differentiation from that match (19). The three lowest % ID values belong to two *Rhodococcus* strains (#135: 71.99% ID, low nutrient, heat-treat, 4°C; and #130: 79.22% ID, low nutrient, heat-treat, 25°C) and one *Bacillus* strain (#46: 78.93% ID, low nutrient, stamp, 25°C).

The third map depicts the availability of recovery nutrients. More numerous and diverse taxonomy was recovered using low-nutrient media (72 isolates, 22 genera). Interestingly, several genera were recovered only within the low-nutrient conditions (*Alcanivorax, Enterobacter, Kocuria, Kytococcus, Micrococcus, Mycobacterium, Pseudomonas, Rossellomorea,* and *Tomitella).* Contrastingly, high-nutrient media resulted in less numerous and less diverse taxonomy (61 isolates, 19 genera); genera solely attributed to high-nutrient recoveries include *Aliihoeflea, Cellulomonas, Nocardia, Nocardioides, Paenibacillus,* and *Pseudoalteromonas*. High-nutrient media selects for faster-growing organisms at the expense of slow-growing organisms (20); the faster-growth equates to competition for the same nutrients required by the less competitive, slow-growing isolates. While this may result in over-representation of certain genera more than others (i.e., *Bacillus*), balance and more diverse bacterial recovery can be achieved through stepwise utilization of selective inoculation and incubation parameters.

To this end of providing selectivity, the fourth map summarizes the inoculation methodology, with the majority of isolates and genera recovered following the more-permissive stamp inoculation (100 isolates, 24 genera) as compared to the restrictive heat-treatment (33 isolates, 11 genera). Inoculation via sediment stamping maintains bacterial associations with both the natural growth substrates (sediment particles) and the community interactions (quorum sensing, metabolic intermediates, etc.) required for recovery of more inter-dependent and potentially rarer taxa (21). Notably, ten isolates with 16S BLASTn % ID < 95% were recovered following sediment-stamping, with five of those isolates (46-*Bacillus,* 219-*Pseudorhizobium,* 136-*Rhodococcus*, 121-*Pseudorhizobium,* and 175-*Rhodococcus*) demonstrating < 90% ID to their closest 16S rRNA sequence NCBI deposits. Contrastingly, heat-treatment decreases population density to only those species able to form heat-resistant spores, or otherwise withstand elevated temperatures for 10 minutes, as demonstrated by the recoveries of the *Mycobacterium* and *Nocardia* isolates. By decreasing the inoculation density (and therefore competition), the remaining population has a greater probability of acquiring nutrients and forming recoverable, viable colonies for sub-passaging, isolation, and identification. Within this reduced population, six isolates demonstrated 16S BLASTn % IDs < 95%, including the 135-*Rhodococcus* isolate with the lowest % ID of 71.991 in comparison to its closest NCBI sequence deposit.

The final map summarizes the initial/recovery incubation temperature. Room temperature/25°C incubation is associated with the recovery of the majority of Gulf of Mexico sediment bacteria (92 isolates, 20 genera). However, despite accounting for less than half the number of room temperature-associated recoveries, the 4°C recovered diversity contains only two less genera (41 isolates, 18 genera). While bacterial recovery results from any combinations of the techniques thus far, it is notable that the colder temperature is associated with a slightly larger proportion of potentially rare isolates (5/41 isolates, ∼12.20%) as compared to the room temperature condition (11/92 isolates, ∼11.96%), as determined by 16S BLASTn % ID < 95%.

### High-nutrient media recovers diverse Gulf of Mexico taxonomy at the expense of quantity

One issue impeding controlled laboratory investigation of biosynthetically interesting organisms is the successful recovery and sustained growth of these environmentally-conditioned (and thusly laboratory-disadvantaged) isolates. To this end of pursuing and characterizing novel chemotaxonomy, we investigated the effects that nutrient availability, inoculation technique, and incubation temperature had upon Gulf of Mexico sediment-associated bacterial biodiversity.

Upon closer inspection of the overall Gulf of Mexico sediment biodiversity, we observed differential recovery of taxonomy potentially correlated with nutritional availability. We employed high-nutrient recovery media as a non-fastidious nutritional comparison to examine the effects that inoculation methods and incubation temperatures had on bacterial recovery. Collectively, using high-nutrient recovery resulted in identification of 61 isolates representing 19 genera within 13 families and 7 taxonomic orders (**Figure 2A**).

**Figure 2.**
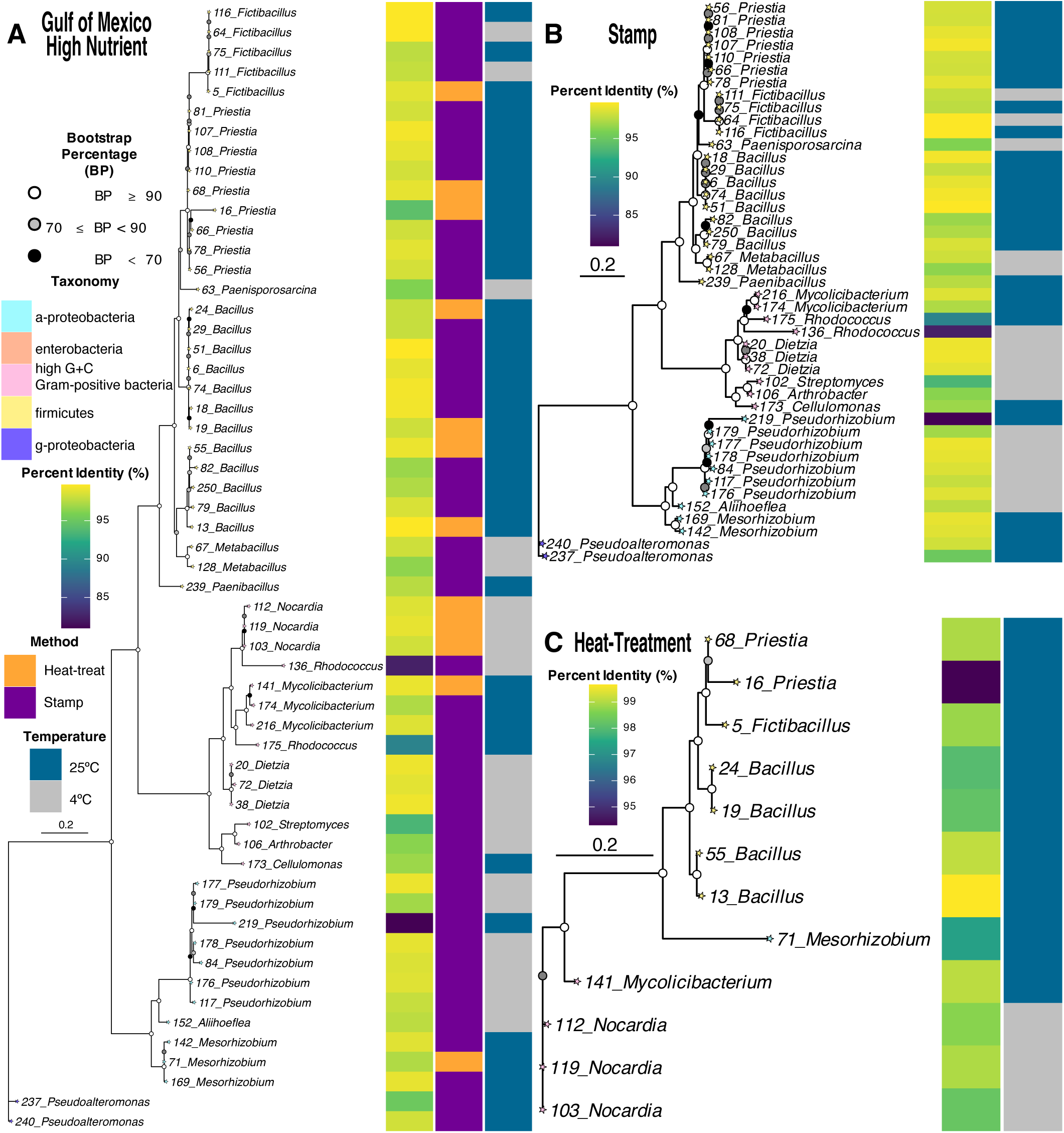
High-nutrient media conditions recover 57 isolates. Shown are all three Gulf of Mexico high nutrient media phylogenetic trees containing most similar genera identifications determined via 16S_ribosomal_RNA BLASTn Database. (**A**) The fifty-seven high nutrient isolates are presented with their BLASTn percent identity values, recovery inoculation method, and recovery incubation temperature. (**B**) The forty-five stamp inoculated and (**C**) twelve heat-treatment inoculated isolates are presented within their respective trees.

In order to examine the impact on biodiversity of the inoculation approach, we contrasted the more permissive sediment stamping inoculation method (**Figure 2B**) with the heat-treated inoculation method (**Figure 2C**) for both room temperature and colder incubations. High nutrient recovery conditions paired with sediment stamping resulted in recovery of forty-eight isolates (18 genera). The remaining high nutrient-associated biodiversity was recovered following the more restrictive heat-treatment (thirteen isolates, 7 genera).

Reviewing the biodiversity of both inoculation techniques on high nutrient media, more isolates and genera were recovered following incubation at room temperature. Notably, despite the risk of fast-growing microbial species overgrowing the high nutrient recovery plates, no fast-growing *Bacillus* spp. were recovered following colder incubations (**Figure 2**). In the absence of rapidly-growing *Bacillus,* rarer and slower-growing biodiversity was observed. Indeed, the only *Nocardia* isolates recovered from the Gulf of Mexico samples belonged to the high nutrient, heat-treated, colder incubation conditions. In addition to growing notoriously slow (22), *Nocardia* have also been characterized as heat-resistant (23). The combination of high nutrient availability, heat-treatment, and colder incubation enabled the recoveries of biodiversity otherwise not recovered within other conditions from Gulf of Mexico samples.

Across our recovery conditions, the rapidly growing *Bacillus* genus persists; interestingly, however, the phylogenetic tree and individual BLASTn’s of the 16S-ITS-23S rRNA gene sequences reveal that these isolates span the spectrum of closely-related and previously-designated *Bacillus* taxonomy. In addition to recovery of closely related *Fictibacillus* (24), *Metabacillus* (25), and *Paenibacillus* (26), we observe recovery of *Priestia* (27). Notably, the #16-*Priestia* isolate has a 16S-ITS-23S rRNA gene sequence percent identity of 94.311% to the closest BLASTn result, indicating that it is within the *Bacillus*-adjacent genera still undergoing taxonomic rearrangement.

### Low nutrient media recovers more numerous isolates

To further characterize culturable Gulf of Mexico sediment-associated bacteria, we employed the same sample processing and recovery techniques on an identical – albeit less concentrated – medium (**Figure 3**). Overall, lower-nutrient media recovered more numerous and diverse taxonomy compared to higher-nutrient conditions (low nutrient: 72 isolates, 22 genera; high nutrient: 61 isolates, 19 genera). Furthermore, nine Gulf of Mexico genera were unique only to low nutrient recovery conditions (*Alcanivorax, Enterobacter, Kocuria, Kytococcus, Micrococcus, Mycobacterium, Pseudomonas, Rossellomorea,* and *Tomitella*).

**Figure 3.**
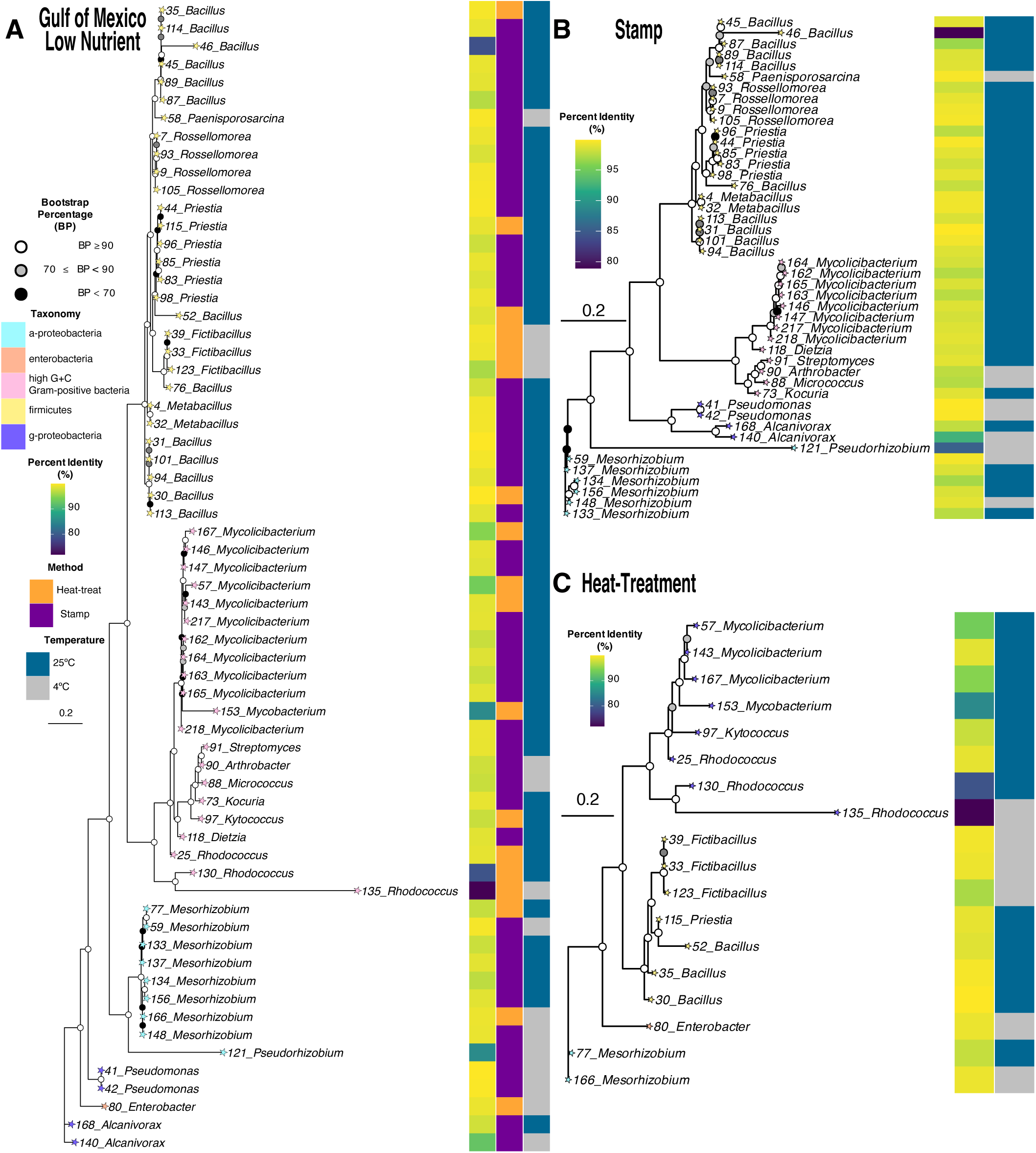
Low-nutrient media conditions recover 64 isolates. Shown are all three Gulf of Mexico low nutrient media phylogenetic trees containing most similar genera identifications determined via 16S_ribosomal_RNA BLASTn Database. (**A**) The sixty-four low nutrient isolates are presented with their BLASTn percent identity values, recovery inoculation method, and recovery incubation temperature. (**B**) The forty-six stamp inoculated and (**C**) eighteen heat-treatment inoculated isolates are presented within their respective trees.

Within the **Figure 2A** overall biodiversity, the 16S-ITS-23S rRNA gene sequence BLASTn percent ID values of isolates to the closest NCBI 16S rRNA database match range from 71.991 – 99.892 percent, with the lower values indicating recovery of rarer and potentially understudied taxonomy. Within this diversity, ten isolates have percent identities <95%, indicating potential taxonomic interest and genera differentiation from the closest NCBI-deposited 16S rRNA gene sequence. Furthermore, this rarer culturable taxonomy includes five isolates with percent identities <86% similarity: #135-*Rhodococcus* (71.991% ID, heat-treat, 4°C), #46-*Bacillus* (78.925% ID, stamp, 25°C), #130-*Rhodococcus* (79.218% ID, heat-treat, 25°C), #153-*Mycobacterium* (85.228% ID, heat-treat, 25°C), and #121-*Pseudorhizobium* (85.448% ID, stamp, 4°C).

Within this taxonomically promising diversity, however, we observe the presence of the highly polyphyletic *Bacillus* and *Bacillus*-adjacent taxa spanning the phylogenetic spectrum within **Figure 3A**. Visually, this polyphyletic clade clusters separately from the other isolates, however, contains multiple sub-clusters that span the continuously refined *Bacillus-/-*like taxonomy. Despite low-GC Gram-positive organisms accounting for ∼43% (31/72 isolates) of low nutrient biodiversity, all isolates were identified as *Bacillus* or genera previously designated as *Bacillus*. Examination of the individual BLASTn searches revealed that these isolates belong to in-flux taxa distinct from the umbrella-like *Bacillus* genus. Of these isolates, three organisms demonstrate BLASTn percent identities <97% that indicate potential species distinction from the closest NCBI-deposited match (#46-*Bacillus*, #123-*Fictibacillus*, #87-*Bacillus)*. In addition to presenting the lowest BLASTn PID, isolate #46-*Bacillus* 16S-ITS-23S rRNA sequence PID of 78.925% indicates the continued need for *Bacillus* taxonomic reclassification. Unsurprisingly, the majority of these fast-growing isolates were recovered following the non-selective sediment stamping technique (**Figure 3B**).

To attenuate potential growth rate-based competition, however, and increase recovery of slower growing taxonomy, we employed a heat-treatment selecting for organisms able to withstand an initial heat shock of 55°C (**Figure 3C**). When examining the heat-treated isolates, we report several taxa that do not form endospores. Although *Bacillus* and *Fictibacillus* are known to produce heat-resistant spores, the remaining isolates are not characterized as such. Instead, these non-spore forming bacteria are known for environmental stress tolerance. *Mesorhizobium* (28), *Mycolicibacterium* (29), and *Rhodococcus* (30) all have literature documenting their ability to overcome environmental stress and elevated temperatures. Interestingly, within this heat-resistant collection, two *Rhodococcus* isolates and two *Mycolicibacterium* isolates demonstrate BLASTn 16S-ITS-23S rRNA sequence PIDs <95%, indicating that they could belong to understudied or novel genera.

Lastly, when examining incubation temperature effects with respect to the two inoculation methods, we again observe that the majority of isolates were recovered at room temperature (**Figures 3B** and **3C**). Notably, however, within that diversity *Arthrobacter, Micrococcus, Paenisporosarcina, Pseudomonas, Rhizobium,* and *Fictibacillus* were only recovered following initial colder incubation temperatures. While their recovery could be resultant of any number of factors, colder incubation exclusivity reveals that culture-dependent biodiversity is improved by including multiple, stepwise incubation parameters. To the end of recovering rare biodiversity, four isolates present 16S-ITS-23S rRNA percent identities <97%, with three isolates demonstrating potential genera divergencies at percent identities <93% (#135-*Rhodococcus* 71.991% ID; #121-*Pseudorhizobium* 85.448% ID; and #140-*Alcanivorax* 92.479% ID).

### A holistic approach to culturing Antarctic sediment-associated bacteria recovers 165 isolates from 3 phyla

When Antarctic sediment samples collected from two depths (20 feet and 60 feet) were examined for culture-dependent biodiversity, the overall complexity of recovered organisms was less than that of the Gulf of Mexico samples. Within **Supplemental Table S3** Antarctic taxonomy, twenty-one genera and ten families are presented. The 16S-ITS-23S rRNA BLASTn results for the Antarctic sediment-associated isolates are presented in **Supplemental Table S4.** These Antarctic isolates are summarized within **Figure 4**, along with their associated BLASTn results, taxonomy, and recovery techniques.

**Figure 4.**
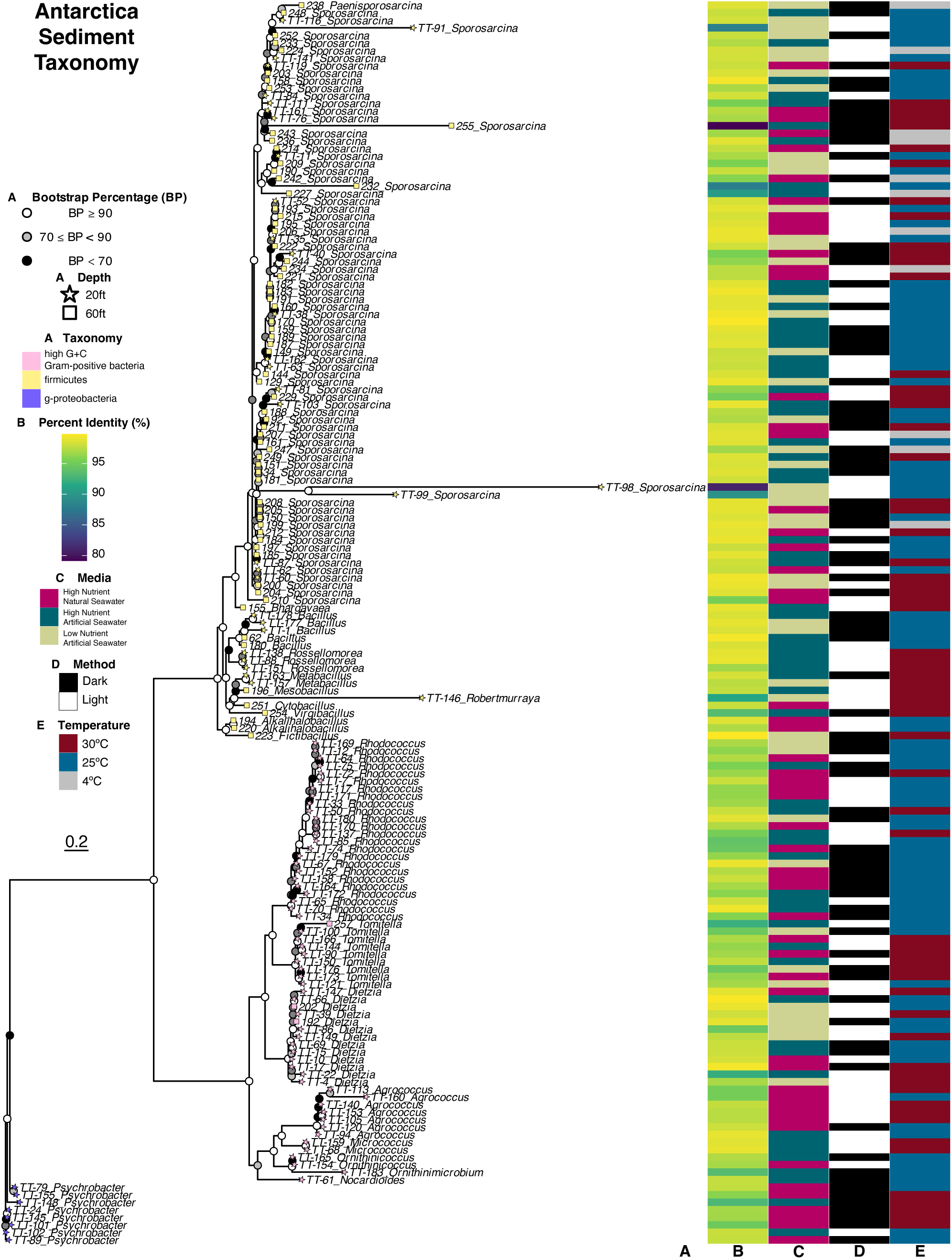
Phylogeny of recovered Antarctic sediment-associated bacteria shows varied taxa represented. The 16S-23S-rRNA sequences from 165 Antarctic isolates were compared to the 16S_ribosomal_RNA BLASTn Database for most similar genera identifications, and then aligned and plotted within the phylogenetic tree (**A**). Isolates’ data is presented as in-laid maps, from left to right: BLASTn percent identity to the most similar result (**B**), recovery media nutrient availability (**C**), recovery light availability (**D**), and recovery incubation temperature (**E**).

Overall, our culture-dependent recoveries are predominantly Actinobacteria and Firmicutes. Despite the relative inaccessibility of Antarctic sediments limiting investigations, several sequencing-dependent studies identify bacterial abundances of Bacteroidetes, Gamma-, Alphaproteobacteria (18), and other Proteobacteria (31). While we demonstrate recovery of these metagenomically-identified taxa, we also recover rarer community members that could otherwise be excluded from culture-independent resolution (32), as these rarer species can be overlooked as a result of primer bias (33). These include the *Agrococcus* (7 isolates)*, Fictibacillus* (1 isolate), and *Psychrobacter* (8 isolates) genera, which have notably escaped culture-independent identifications (32).

To optimize recovery of this under-investigated Antarctic biodiversity, we modified our Gulf of Mexico culture techniques to attenuate the culture barrier facing Antarctic strains. Instead of employing the heat-treatment processing used on the Gulf of Mexico isolates, we chose to only use the permissive sediment-stamping inoculation. To further attenuate the culture barrier, we extended the initial incubation, as previous literature reports that extended incubations are successful in recovering otherwise recalcitrant biodiversity (34). Additionally, we examined the potential impact of natural versus artificial seawater within recovery media, as there have been reports of obligatory seawater organisms (35). Lastly, we included variations in light-availability during initial incubations; while there are well-documented circadian clocks within photosynthetic organisms, there is also growing evidence for non-photosynthetic bacterial circadian rhythms (36).

Within the overall diversity, fifty-one isolates contain 16S-ITS-23S rRNA percent identities <97% to their closest NCBI-deposited match, indicating potential species differentiation. Notably, eighteen of those isolates’ percent identities fall below the <95% genera differentiation threshold and could present understudied sources for novel chemotaxonomy. These isolates belong to eight genera: *Sporosarcina, Robertmurraya, Dietzia, Tomitella, Psychrobacter, Virgibacillus, Ornithinimicrobium,* and *Rhodococcus.* To investigate rarer sediment bacteria, we examined the recovered biodiversity of both the 20-foot and 60-foot depth samples within their respective sampling depths.

### The majority of Antarctic sediment-associated bacteria were recovered from samples collected at 20-foot depths

Ninety-five of the 165 isolates (∼58%) were recovered from the sediment samples collected from 20-foot depths. A total of 14 genera were represented within the 20-foot depth samples (**Figure 5A**). Notably – despite their *Bacillus-*like expedited growth rates, abilities, and wide environmental distribution – the only *Metabacillus* (25)*, Robertmurraya* and *Rossellomorea* (27) Antarctic isolates were limited to the 20-foot depth sediments. These genera – in addition to many others – have only been distinguished from the diverse *Bacillus* genus recently, indicating that their increasing environmental recoveries could continue to guide taxonomic rearrangement.

**Figure 5.**
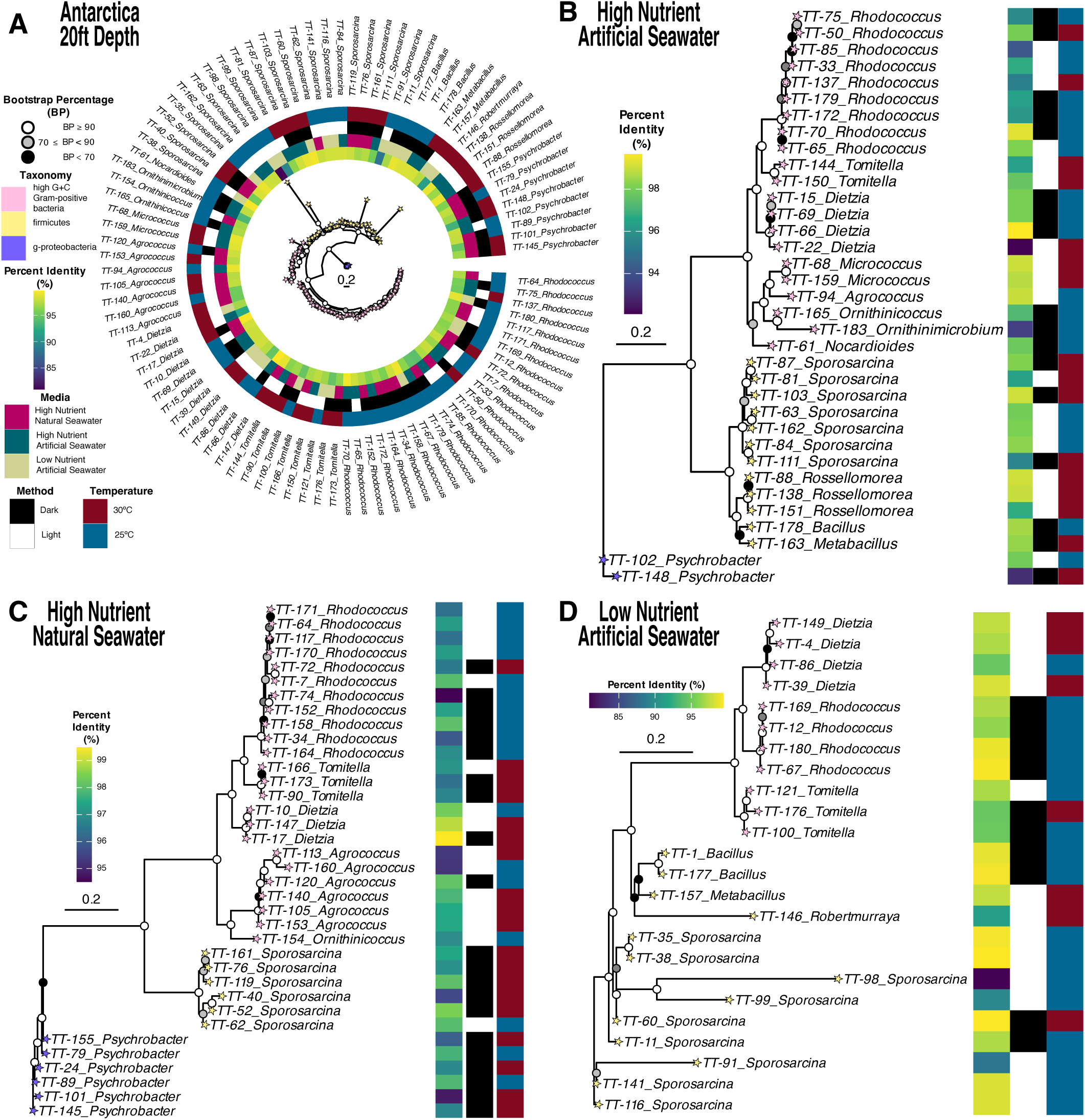
Varied recovery techniques result in isolation of 95 isolates from Antarctic sediment collected from 20ft depths. All four Antarctic 20-ft depth phylogenetic trees contain the most similar genera identifications determined via 16S_ribosomal_RNA BLASTn Database. (**A**) The ninety-five Antarctic 20-ft depth isolates are presented with their BLASTn percent identity values, recovery media nutrient availability, recovery light availability, and recovery incubation temperature. (**B**) The thirty-five high nutrient + artificial seawater isolates, (**C**) thirty-six high nutrient + natural seawater, and (**D**) twenty-four low nutrient + artificial seawater isolates are presented within their respective trees.

To the end of increasing environmental recovery of taxonomically interesting bacteria, we first examined the effects of nutrient-availability on recovery. Although both the high-(**Figures 5B** – **5C**) and low-nutrient (**Figure 5D**) conditions resulted in similar biodiversity, the only Antarctic isolate unique to low-nutrient conditions was the sole *Robertmurraya* species. Since the genus was established in 2020, *Robertmurraya* currently consists of seven validly-published species according to the List of Prokaryotic names with Standing in Nomenclature (LPSN, https://lpsn.dsmz.de/genus/robertmurraya). Although *Robertmurraya* type species have been isolated from soil (37, 38, 39), biowaste composting reactors (40), silage (41), and fecal flora (42), our recovered isolate has the highest sequence similarity to the psychrotolerant *Robertmurraya beringensis* seawater isolate (**Supplemental Table S4**; (43)).

To further examine nutrient-availability, we examined the high-nutrient conditions for isolates only recoverable on natural seawater-based media (**Figure 5C**). While the recovery numbers were similar regardless of seawater type (36 isolates in natural seawater; 35 isolates in artificial seawater), the artificial seawater conditions recovered six more unique genera than the natural seawater conditions (*Bacillus, Metabacillus, Micrococcus, Nocardioides, Ornithinimicrobium,* and *Rossellomorea*).

We next examined light availability and temperature during initial incubations and observed impacts on recovery. While light-restricted incubations recovered similar biodiversity as freely light-available conditions, the only Antarctic species unique to darkened incubations belonged to the *Bacillus, Nocardioides,* and *Ornithinimicrobium* genera (**Figure 5**). Although not as significant as nutrient- or light-availability on recovered biodiversity, we did observe temperature-specific recovered biodiversity. Across all nutrient and light conditions, the majority of the 20-foot depth isolates were recovered following room temperature incubation (53/95 isolates, ∼56%, **Figure 5**). Contrastingly, the only genera unique to the 30°C incubation were the *Metabacillus, Micrococcus, Robertmurraya,* and *Rossellomorea* isolates.

Overall, we observe that a holistic approach to culture conditions resulted in a unique, culture-dependent insight to Antarctic biodiversity. Without employing the entirety of the tested conditions, several chemotaxonomically interesting isolates would have otherwise remained unidentified. This holistic approach is most apparent when examining the nutrient conditions’ proportions of potentially rare taxonomy (where 16S-ITS-23S rRNA percent identities <95% to their closest NCBI-deposited match): as a whole, the low-nutrient media (7/24 isolates, 29.17%) resulted in the greatest proportion of potentially rare isolates. Notably, the use of artificial or natural seawater resulted in a disproportionate effect on potentially rare biodiversity, where a greater proportion of potentially rare isolates were recovered on artificial seawater (4/35 isolates, 11.43%) as compared to the same nutrient-media recipe utilizing natural seawater (2/36 isolates, 5.56%).

### Comprehensive culturing approaches result in taxonomically diverse Antarctic bacteria isolated from sediment samples collected at 60-foot depths

Seventy of the 165 Antarctic isolates (∼42%) were recovered from sediment samples collected from 60-foot depths, representing eleven genera (**Figure 6A**). Although we observed decreased biodiversity as compared to the 20-foot depth samples, we did recover seven unique genera not represented within the 20-foot depth samples. These were *Alkalihalobacillus, Bhargavaea, Cytobacillus, Fictibacillus, Mesobacillus, Paenisporosarcina,* and *Virgibacillus.* Notably, the *Alkalihalobacillus, Cytobacillus* and *Mesobacillus* (25), *Fictibacillus* (24), and *Virgibacillus* (44) genera previously belonged to the *Bacillus* taxon. Although bacterial taxonomy can be transiently in-flux, continued recovery of environmental biodiversity can only improve taxonomic identification.

**Figure 6.**
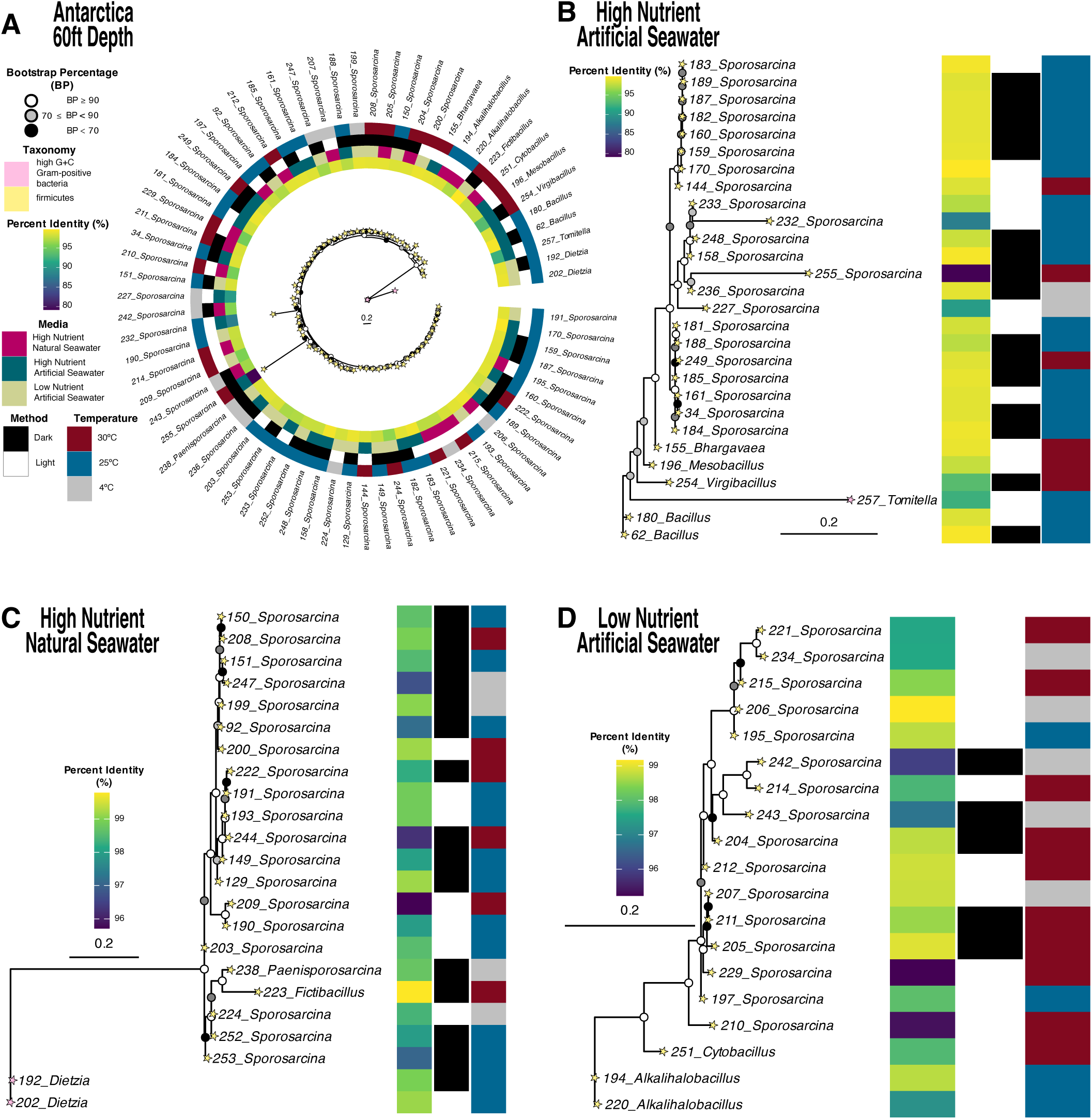
Varied recovery techniques result in isolation of 70 isolates from Antarctic sediment collected from 60ft depths. All four Antarctic 60-ft depth phylogenetic trees contain the most similar genera identifications determined via 16S_ribosomal_RNA BLASTn Database. (**A**) The seventy Antarctic 60-ft depth isolates are presented with their BLASTn percent identity values, recovery media nutrient availability, recovery light availability, and recovery incubation temperature. (**B**) The twenty-eight high nutrient + artificial seawater isolates, (**C**) nineteen high nutrient + natural seawater, and (**D**) twenty-three low nutrient + artificial seawater isolates are presented within their respective trees.

In order to elicit growth of such taxonomically-important biodiversity, we first employed differential nutrient conditions. High-nutrient conditions (**Figures 6B** and **6C**) collectively recovered a greater number and diversity of isolates as compared to low-nutrient conditions (**Figure 6D**). Although *Sporosarcina* isolates were recovered in all three nutrient conditions, the recovery of the remaining 60-foot depth-associated genera correlated with only one nutrient condition. Notably, when examining the effects of seawater (natural and artificial), we observed the recovery of three isolates unique to the natural seawater condition: two *Alkalihalobacillus* isolates and one *Cytobacillus.* The etymology of *Alkalihalobacillus* correctly implies that the rod-shaped bacteria can survive in both salty and alkaline conditions; this is supported by the recovery only following incubation with natural seawater.

To further provide a more comprehensive culture-dependent examination of Antarctic biodiversity, we also examined the effects of temperature and light-availability (**Figure 6**). The majority of isolates were recovered following incubation at room temperature, followed by 30°C and 4°C. Although Antarctic isolates were anticipated to prefer colder conditions, we only recovered 11/70 (∼16%) with colder incubation temperatures. Notably, ten isolates belonged to the *Sporosarcina* genus, with the remaining isolate belonging to the closely related and evolutionarily divergent *Paenisporosarcina* genus. Similarly to *Bacillus* taxonomic reclassification, *Sporosarcina* has undergone rearrangement to produce a distinct *Paenisporosarcina* clade (45). Although multiple conditions contributed to recovery of *Paenisporosarcina*, it is notable that the most restrictive conditions (light-restricted and low-nutrient and temperature) facilitated its recovery, rather than constricted its viability.

### Multifactorial analysis reveals significant impacts on recovered bacterial taxonomy within sediment samples

Collectively, 286 isolates within 38 genera were recovered from both collection sites. Notably, with the exception of eight shared genera (*Bacillus, Dietzia, Fictibacillus, Metabacillus, Micrococcus, Paenisporosarcina, Rhodococcus,* and *Rossellomorea*), the two sampling sites resulted in geographically-distinct recoverable biodiversities (**Figure 7**). Gulf of Mexico recovered biodiversity is predominately *Bacillus, Mycolicibacterium,* and *Priestia* (**Figure 7A**); notably, all isolates within these genera were recovered after initial incubation at 25°C. Interestingly, while *Bacillus* was recovered from both regions, Antarctic *Bacillus* recovery is less numerous, and recovery is similarly restricted to incubation at 25°C (**Figure 7B**). Antarctic recovered biodiversity is predominately *Rhodococcus* and *Sporosarcina* (**Figure 7B**). While *Rhodococcus* isolates were solely recovered from 20ft-depth Antarctic sediment samples, *Sporosarcina* isolates were predominately associated with 60ft-depth samples.

**Figure 7.**
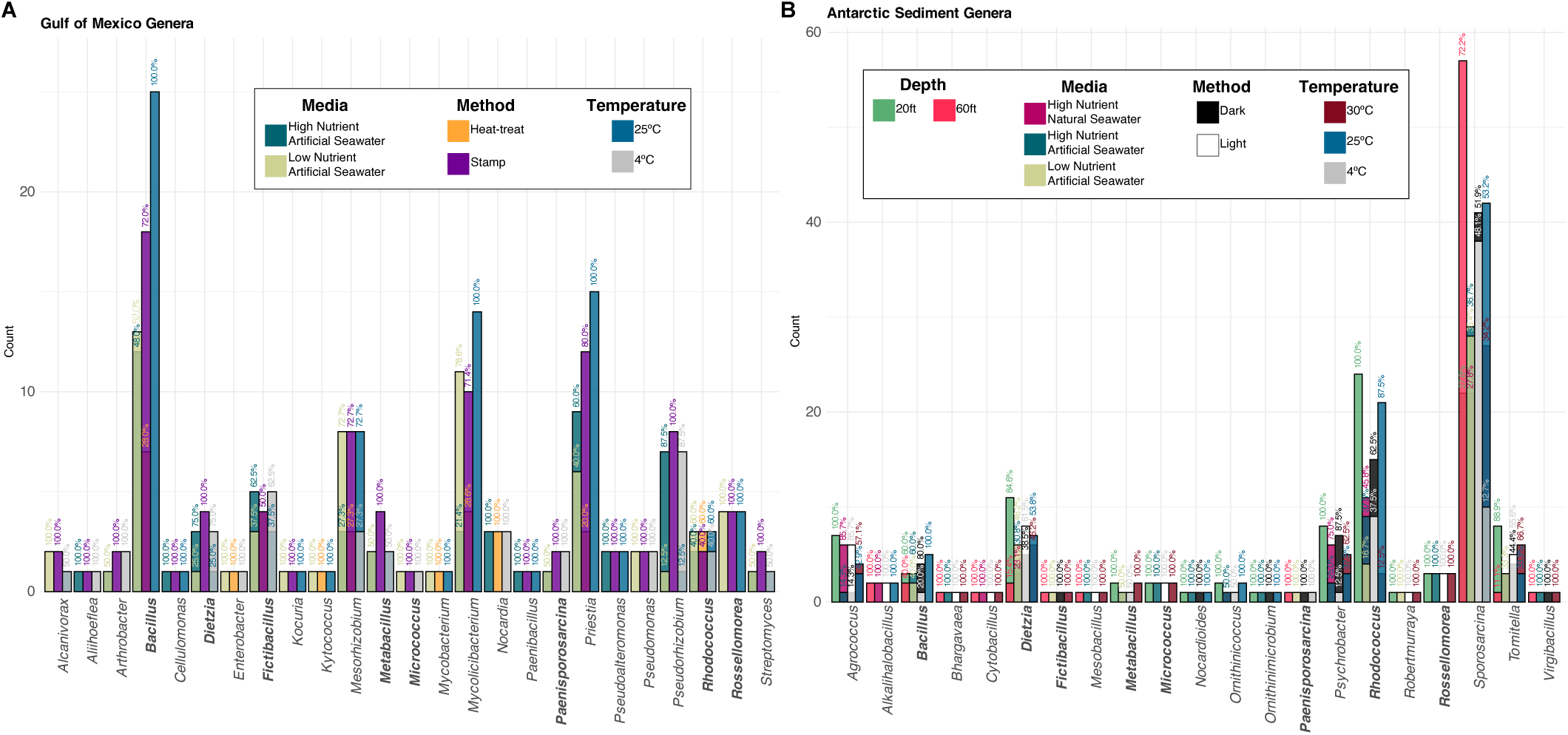
Total recovered sediment-associated cultivars from Gulf of Mexico and Antarctica. Isolate counts and associated recovery data within recovered genera are presented for the Gulf of Mexico (**A**) and Antarctic (**B**) sediment-associated cultivars. Media types, inoculation methodologies, incubation temperatures, and isolation frequencies are plotted for both sampling sites, with sediment sampling depths specific to Antarctic genera.

While visual trends can be interpreted from such an overview of recovery, multinomial logistic regression can begin to explore how these different recovery conditions influence the isolation of bacterial genera. We modeled the relationship between recovery of Gulf of Mexico genera and culture conditions, considering the baseline/reference comparison as the recovery combination associated with the largest number of recovered isolates, or an otherwise “generalist” approach (Isolate: *Bacillus*; Media: Low Nutrient, Artificial Seawater; Method: Stamp; Temperature: 25°C). Although our dataset is constrained by the infrequency of culturable genera, the majority of the genera demonstrate significant recovery preferences (**Supplemental Table S5**) compared to the generalized approach used to recover the greatest number of isolates. Indeed, by examining the overall recovery odds if all conditions were chosen to favor larger recovery numbers (generalization), we determine that the majority of genera present decreased likelihoods of recovery. While recovery is undoubtedly resultant of multiplicative factors, examining individual conditions yielded numerous insights into the recoverable biodiversity. Although low nutrient media resulted in the largest number of recovered isolates and preference towards certain genera (*Rossellomorea, Pseudomonas,* and *Alcanivorax*), several genera present with higher odds of recovery within high nutrient media (*Pseudoalteromonas, Cellulomonas,* and *Paenibacillus*). While nutrient availability appears to be statistically impactful on recovery (log odds ranging in magnitude from −33 to 40; average −2; median −0.8), the inoculation methodology presents with more genera demonstrating similarly large magnitudes of impact (ranging from −30 to 34; average −9; median −15). In addition to recovering the majority of isolates, the stamp inoculation technique is associated with higher odds of recovery of more unique genera as compared to the heat-treatment inoculation. Stamp-inoculation results in higher recovery odds of *Pseudorhizobium, Pseudomonas, Rossellomorea, Arthrobacter, Paenisporosarcina,* and *Alcanivorax;* contrastingly, heat-treatment favors recoveries of *Kytococcus* and *Mycobacterium.* Initial incubation at 25°C favored *Mycolicibacterium, Rossellomorea, Priestia,* and *Pseudoalteromonas;* however, decreased temperatures (4°C) favored recovery of *Pseudomonas, Arthrobacter,* and *Paenisporosarcina.* The log odds ranging from −40 to 59 (average 18; median 26) indicate that, although the majority of isolates were recovered following the increased 25°C incubation, numerous genera demonstrate stronger odds of recovery following decreased/4°C incubations.

We next modeled the Antarctic recovery data for insights into recovery preferences, focusing on sediment sample depth, nutrient and light availability, and initial incubation temperature (**Supplemental Table S6**). The baseline/reference comparison for Antarctic recovery was the combination resulting in the largest number of recovered bacteria (Isolate: *Sporosarcina*; Depth: 60ft; Media: High Nutrient, Artificial Seawater; Method: Dark; Temperature: 25°C). Although this combination resulted in the most numerous recoveries, the overall analysis indicated increased odds of recovery associated with a more individualized approach (generalized approach recovery odds range from −437 to −2; average −230; median −240). Recovery odds were greatly impacted by originating sediment sample depth. Although the majority of isolates were isolated from 20ft-deep sediment samples, the most generalized recovery combination was associated with 60ft-deep samples (recovery odds ranging −88 to 325; average 83; median 96). As such, more genera demonstrated higher odds of recovery from 20ft depths (including *Agrococcus, Rhodococcus,* and *Psychrobacter*) as compared to those strongly-associated with the deeper 60ft samples (*Sporosarcina* and *Alkalihalobacillus*). Although two of the examined nutrient conditions contained the same composition, they differed on the seawater type (artificial or natural) used to solvate ingredients prior to sterilization and subsequent aseptic inoculation of sediment sample bacteria. Overall, the majority of Antarctic isolates were recovered with high nutrient media, with the most generalized (and frequent) recovery associated with artificial seawater (odds ranging −169 to 160; average −43; median −17). Notably, several high-nutrient associated genera demonstrate higher recovery odds associated with artificial seawater (*Rossellomorea* and *Micrococcus*) as compared to those more likely to be recovered with natural seawater (*Alkalihalobacillus*). Contrastingly, low nutrient artificial seawater recovery odds were overall lower in comparison to high nutrient recovery (odds ranging −147 to 115; average −48; median −60). Interestingly, although *Dietzia* recovery was possible with all tested media, recovery odds were highest when associated with low nutrient conditions. Initial incubations within dark conditions resulted in the most numerous recoveries, however, several genera demonstrated preferences for light-available conditions (*Rossellomorea* and *Micrococcus;* overall odds ranging −82 to 149; average 24; median 0.1). Room temperature (25°C) initial incubation resulted in the greatest number of recoveries, however, several taxa demonstrate increased recovery odds following incubation at 30°C (*Metabacillus, Rossellomorea,* and *Micrococcus*; overall odds ranging −152 to 164; average 27; median 19). Notably, however, the predictive model does not concur with the observed recovery preferences for the initial 4°C incubation. Despite several taxa demonstrating 4°C incubation preferences, the log odds indicate very low likelihoods of successful recovery (*Bacillus*, −108; *Rhodococcus* −94; and *Ornithinicoccus* −63).

## Concluding Remarks

Our holistic and culture-dependent approach to Gulf of Mexico and Antarctic sediment samples presents unique insights into prokaryotic biodiversity often overlooked by prevailing culture-independent surveys. In addition to fostering laboratory sustainable cultivars of potentially low-abundant/rarer taxa, our culture-dependent findings enable empirical investigations beyond those solely restricted to *in silico* studies.

To this end, in addition to recovering key members from traditionally sediment-associated taxa (i.e., Pseudomonadota phyla) (46), our techniques have also recovered a substantial number of unique taxa within the Actinomycetota phylum (35/121 Gulf of Mexico isolates, ∼29%; and 59/165 Antarctic isolates, ∼36%). Critically, while this phylum serves as one of the most prolific (and continuous) sources of natural products (47), the majority of available metagenomic datasets do not report Actinomycetota in as high of an abundance as that found within our samples, indicating the efficacy of our approach to recovering otherwise under-represented taxa. Additionally, our not insignificant Antarctic Actinobacterial fraction (59/165 Antarctic isolates, ∼36%) is notable because, in addition to the taxon’s innumerable bioactive compounds, Antarctic bacteria have evolved survival mechanisms to overcome unique environmental stressors like extreme cold, UV radiation, and salinity (48). Subsequently, these evolved mechanisms can be capitalized for bioremediation, biotechnology, and drug discovery efforts. Thorough investigations of this much sought-after biosynthetic potential, however, is only possible following the transition of environmental microbe to laboratory-viable organism.

Additionally, our culture-dependent efforts provide ecological insights into the sampled environments. Our Gulf of Mexico sediment sample biodiversity contains key Deepwater Horizon taxa (Gammaproteobacteria, Alphaproteobacteria, and Actinomycetes (49, 50, 51)) and other genera associated with hydrocarbon-degrading activities (*Alcanivorax, Rhodococcus,* and *Pseudomonas*). While bacterial consortia are increasingly recognized as responsible for the final and complete detoxification of oil-contaminated marine environments (52), the most abundant taxon is almost always *Alcanivorax,* accounting for >80% of the oil-associated populations (53), indicating its bioremediation potential (54).

Overall, our chemotaxonomically interesting sediment-associated isolates (both within the Gulf of Mexico and Antarctic) are otherwise under-reported within their respective sites’ high-throughput metagenomic sequencing. Although our cultivars’ sequences are similar to those organisms demonstrating chemotaxonomic promise, the lack of culture-independent precedence for our recovered biodiversity is not unique. While no single explanation can fully resolve the discrepant biodiversity estimations, several contributing factors have been identified. Prior to complete degradation, nonviable/dead environmental DNA can be detected within culture-independent studies (55, 56), thereby overinflating a sample’s expected biodiversity. Furthermore, these primarily sequence-identified bacteria may belong to viable but nonculturable (VBNC) taxa that could enter a dormant/non-replicative state within the laboratory (55), again, producing an *in silico* overestimation of an environment’s *accessible* biodiversity. Lastly, sequencing biases (including primer design (57, 58) and amplicon sequencing depth (59)) may overlook rarer bacterial diversities that would otherwise remain unidentified without culture-dependent efforts.

Collectively, our culturing approach to Gulf of Mexico and Antarctic sediment samples has succeeded in recovering 286 laboratory-viable cultivars. Although culture-independent investigations of the originating/proximal sampling site may include additional/non-recovered taxa, our approach has expanded methodologies to enhance bacterial recovery. By expanding the culturing conditions (nutrition, temperature, light availability, etc.), we recover a larger/richer biodiversity from environmental sediment samples, as similarly observed in other marine sediment samples (60) and the human gut (61). This laboratory-viable environmental bacterial collection and their recovery methodologies can be extrapolated to similar taxa and geographies, potentially enabling future investigations into increasingly rarer bacterial diversity.

## Experimental Procedures

### Environmental sample collection and processing

The Gulf of Mexico sediment samples were collected from the Clearwater Reef and Florida Middle Grounds areas in 2015 via scuba diving and stored at room temperature for 2 years prior to processing. Antarctic sediment was collected from East Arthur Harbor, Palmer Station in 2018 from 20-foot and 60-foot depths. The samples were stored at −20°C in the collected seawater. All sediment samples were processed following the methods of Jensen PR et al (62). Briefly, ∼5 g of the wet sediment was dried overnight in a sterile laminar flow hood. An autoclaved artificial sponge was pressed into the dried sediment and then stamped onto the surface of an agar plate in a clockwise, serial dilution manner. This was repeated for multiple nutrient and incubation conditions (see below). The remaining dried Gulf of Mexico sediment, only, was then added to ∼3 mL of sterile artificial seawater, Instant Ocean® (IO; 36 g/L deionized water), and vigorously vortexed. The mixture was then heated in a water bath at 55°C for 10 minutes before plating 75 μL onto agar.

### Bacterial recovery media and incubation

Two nutritional media were used to recover the Gulf of Mexico bacterial-associated sediment samples. The higher-nutrient media, AMM (62), contains 18 g agar, 10 g starch, 4 g yeast extract, and 2 g peptone for 1L of IO water. The lower-nutrient media, M1low (63), contains 18 g agar, 2 g starch, 0.8 g yeast extract, and 0.4 g peptone for 1L of IO water. Gulf of Mexico samples were incubated at room temperature (∼25°C) and 4°C for at least 4 months.

The Antarctic sediment samples were recovered using AMM and M1low (both as described above), and one additional media type: A1 (64), which contains the same composition as AMM, however, replaces the artificial seawater with naturally obtained seawater from the Gulf of Mexico. The natural seawater is autoclaved during the initial sterilization of the nutrient-agar solutions. To aid in recovery of slower-growing isolates, we also employed the use of dark conditions by wrapping in aluminum foil to obstruct light permeation to the incubated plates. Antarctic samples were incubated at 4°C, room temperature (∼25°C), and 30°C for at least 4 months in both light and dark conditions. All incubation combinations were performed with technical duplicate agar plates within the parallel recovery media tested.

Colony growths were aseptically sub-passaged for isolation on International *Streptomyces* Project medium 2 (10 g malt extract, 4 g glucose, 4 g yeast extract, and 15 g agar for 1L IO water), and incubated at room temperature or 30°C. Upon isolation, strains were grown in 5 mL of tryptic soy broth (TSB; 30 g for 1 L of deionized water) supplemented with three sterile glass beads (∼1 mm diameter) and a final concentration of 20% (v/v) solution of filter-sterilized (0.22 μm-pore size, cellulose acetate filters) 50% (w/v) sucrose for 7-14 days at 210 rpm and 30°C. Upon turbidity, cultures were cryopreserved within 20% (v/v) glycerol-TSB solutions at −80°C.

### DNA extraction, PCR amplification, Sanger sequencing, and BLASTn identification

For ITS-identification, cryopreserved glycerol stocks were streaked on tryptic soy agar (TSA; 30 g TSB and 15 g agar for 1 L of deionized water) supplemented with an overlay of 250 μL of 50% (w/v) sucrose solution. Plates were then incubated at 30°C until visible growth was obtained. Isolated colonies were then picked from the agar plates and grown in 5 mL of tryptic soy broth (TSB; 15g for 1L deionized water) supplemented with sterile glass beads and a final concentration of 20% (v/v) solution of filter sterilized 50% (w/v) sucrose for 7-14 days at 210 rpm and 30°C. Upon obtaining visibly turbid growth, cultures were mechanically disrupted with sterile glass beads (∼0.1 mm) before undergoing phenol – chloroform DNA extraction. Quality was monitored with a Thermo Scientific NanoDrop® ND-1000 UV-Vis Spectrophotometer.

Organism identification via 16S rRNA gene sequencing was performed via PCR using 16S-ITS-23S rRNA primers (F 5′-CCRAMCTGTCTCACGACG-3′ and R 5′-CCRAMCTGTCTCACGACG-3′ (65)). Products were purified using a QIAquick PCR purification Kit (Qiagen) following the manufacturer’s instructions. Sanger sequencing using the above forward primer was performed (Genewiz/Azenta Life Sciences), and sequences were subjected to a NCBI BLASTn (v2.12.0+) search against the 16S_ribosomal_RNA database (downloaded August 7, 2022 at 3:51pm) to ascertain a minimum of genus level identification.

### Phylogenetic trees

The 16S-ITS-23S rRNA Sanger-derived sequences were labeled with the most likely BLASTn-identified genus (results for Gulf of Mexico presented in **Supplemental Table S2** and Antarctica in **Supplemental Table S4**), and aligned via MAFFT v7.505 (2022/Apr/10) (66), enabling the -auto function within each subgroup (location, media, inoculation). Phylogenetic trees based on the aligned 16S-ITS-23S rRNA sequences were generated via PhyML 3.3.20211231 (default settings) (67). The ggTree v3.4.1 (68, 69), ggnewscale v0.4.9 (70), and ggtreeExtra v1.10.0 (71, 72) R packages were used to generate the phylogenetic trees and heatmaps. The ape v5.7-1 (73) package in R was used to calculate branch lengths.

### Multinomial log-linear modeling

R package nnet v7.3-19 (74) was used to predict recovery odds via multinomial log-linear modeling (Gulf of Mexico formula: Isolate ∼ Media + Method + Temp; Antarctica formula: Isolate ∼ Depth + Media + Method + Temp). Two-tailed z-tests were conducted to assess the statistical significance of multinomial logistic regression coefficients.

## Data availability

The NCBI GenBank accession numbers for the 16S-ITS-23S rRNA Sanger-derived sequences are PP778693-PP778978. The statistical script and input data are available at https://github.com/DrSarahJKennedy/culture-condition-preferences.

## Acknowledgments

This study was supported by grants AI124458 and AI157506 (both to L.N.S.) and AI154992 (B.J.B. and L.N.S.) from the National Institute of Allergy and Infectious Diseases.

