## Supplementary Information for "Culture-dependent identification of rare marine sediment bacteria from the Gulf of Mexico and Antarctica"

**Supplemental Table S1.** Recovered Gulf of Mexico sediment-associated taxonomy spans 3 phyla.

| Genera | Family | Order | Class | Phylum |
|---|---|---|---|---|
| *Alcanivorax* | *Alcanivoracaceae* | Oceanospirillales | Gammaproteobacteria | Pseudomonadota |
| *Aliihoeflea* | *Phyllobacteriaceae* | Hyphomicrobiales | Alphaproteobacteria | Pseudomonadota |
| *Arthrobacter* | *Micrococcaceae* | Micrococcales | Actinomycetes | Actinomycetota |
| *Bacillus* | *Bacillaceae* | Caryophanales | Bacilli | Bacillota |
| *Cellulomonas* | *Cellulomonadaceae* | Micrococcales | Actinomycetes | Actinomycetota |
| *Dietzia* | *Dietziaceae* | Mycobacteriales | Actinomycetes | Actinomycetota |
| *Enterobacter* | *Enterobacteriaceae* | Enterobacterales | Gammaproteobacteria | Pseudomonadota |
| *Fictibacillus* | *Bacillaceae* | Caryophanales | Bacilli | Bacillota |
| *Kocuria* | *Micrococcaceae* | Micrococcales | Actinomycetes | Actinomycetota |
| *Kytococcus* | *Kytococcaceae* | Micrococcales | Actinomycetes | Actinomycetota |
| *Mesorhizobium* | *Phyllobacteriaceae* | Hyphomicrobiales | Alphaproteobacteria | Pseudomonadota |
| *Metabacillus* | *Bacillaceae* | Caryophanales | Bacilli | Bacillota |
| *Micrococcus* | *Micrococcaceae* | Micrococcales | Actinomycetes | Actinomycetota |
| *Mycobacterium* | *Mycobacteriaceae* | Mycobacteriales | Actinomycetes | Actinomycetota |
| *Mycolicibacterium* | *Mycobacteriaceae* | Mycobacteriales | Actinomycetes | Actinomycetota |
| *Nocardia* | *Nocardiaceae* | Mycobacteriales | Actinomycetes | Actinomycetota |
| *Paenibacillus* | *Paenibacillaceae* | Caryophanales | Bacilli | Bacillota |
| *Paenisporosarcina* | *Caryophanaceae* | Caryophanales | Bacilli | Bacillota |
| *Priestia* | *Bacillaceae* | Caryophanales | Bacilli | Bacillota |
| *Pseudoalteromonas* | *Pseudoalteromonadaceae* | Alteromonadales | Gammaproteobacteria | Pseudomonadota |
| *Pseudomonas* | *Pseudomonadaceae* | Pseudomonadales | Gammaproteobacteria | Pseudomonadota |
| *Pseudorhizobium* | *Rhizobiaceae* | Hyphomicrobiales | Alphaproteobacteria | Pseudomonadota |
| *Rhodococcus* | *Nocardiaceae* | Mycobacteriales | Actinomycetes | Actinomycetota |
| *Rossellomorea* | *Bacillaceae* | Caryophanales | Bacilli | Bacillota |
| *Streptomyces* | *Streptomycetaceae* | Streptomycetales | Actinomycetes | Actinomycetota |

**Supplemental Table S2.** Gulf of Mexico sediment bacterial isolates' BLASTn results (1/14).

| Isolate | Result_ID | Result_Sci_Names | Subject_Blast_Names | %ID | ALIGN_LEN | Mismatches | Gap Opens | Q. start | Q. end | S. start | S. end | Evalue | Bit Score | % Query Coverage Per Subject |
|---|---|---|---|---|---|---|---|---|---|---|---|---|---|---|
| 5 | gi\|636560458\|ref\|NR_116518.1\| | *Fictibacillus rigui* | firmicutes | 98.817 | 845 | 8 | 2 | 15 | 858 | 541 | 1384 | 0 | 1478 | 97 |
| 13 | gi\|1269801501\|ref\|NR_075005.2\| | *Bacillus velezensis* | firmicutes | 99.66 | 883 | 2 | 1 | 15 | 896 | 567 | 1449 | 0 | 1578 | 98 |
| 16 | gi\|645321408\|ref\|NR_118382.1\| | *Priestia flexa* | firmicutes | 94.311 | 791 | 33 | 10 | 12 | 792 | 538 | 1326 | 0 | 1214 | 89 |
| 19 | gi\|1441204320\|ref\|NR_157735.1\| | *Bacillus proteolyticus* | firmicutes | 98.194 | 886 | 13 | 3 | 20 | 904 | 567 | 1450 | 0 | 1527 | 98 |
| 24 | gi\|1441204320\|ref\|NR_157735.1\| | *Bacillus proteolyticus* | firmicutes | 98.018 | 908 | 13 | 5 | 22 | 925 | 567 | 1473 | 0 | 1551 | 98 |
| 55 | gi\|1212229176\|ref\|NR_148244.1\| | *Bacillus xiamenensis* | firmicutes | 99.122 | 911 | 7 | 1 | 15 | 924 | 561 | 1471 | 0 | 1610 | 98 |
| 68 | gi\|636559893\|ref\|NR_115953.1\| | *Priestia aryabhattai* B8W22 | firmicutes | 98.963 | 964 | 9 | 1 | 13 | 975 | 567 | 1530 | 0 | 1698 | 94 |
| 71 | gi\|1315204046\|ref\|NR_152716.1\| | *Mesorhizobium sediminum* | a-proteobacteria | 97.345 | 791 | 18 | 3 | 12 | 801 | 499 | 1287 | 0 | 1336 | 99 |
| 103 | gi\|559795187\|ref\|NR_104776.1\| | *Nocardia coeliaca* | high G+C Gram-positive bacteria | 98.274 | 985 | 14 | 3 | 13 | 994 | 523 | 1507 | 0 | 1702 | 95 |
| 112 | gi\|559795187\|ref\|NR_104776.1\| | *Nocardia coeliaca* | high G+C Gram-positive bacteria | 98.684 | 988 | 8 | 5 | 14 | 999 | 523 | 1507 | 0 | 1713 | 96 |
| 119 | gi\|559795187\|ref\|NR_104776.1\| | *Nocardia coeliaca* | high G+C Gram-positive bacteria | 98.976 | 977 | 7 | 3 | 15 | 990 | 523 | 1497 | 0 | 1712 | 98 |

**Supplemental Table S2 (continued):** Gulf of Mexico sediment bacterial isolates' BLASTn results (2/14).

| Isolate | Result_ID | Result_Sci_Names | Subject_Blast_Names | %ID | ALIGN_LEN | Mismatches | Gap Opens | Q. start | Q. end | S. start | S. end | Evalue | Bit Score | % Query Coverage Per Subject |
|---|---|---|---|---|---|---|---|---|---|---|---|---|---|---|
| 141 | gi\|559795186\|ref\|NR_104775.1\| | *Mycolicibacterium fortuitum* subsp. *acetamidolyticum* | high G+C Gram-positive bacteria | 99.084 | 983 | 5 | 4 | 17 | 995 | 523 | 1505 | 0 | 1722 | 85 |
| 6 | gi\|1441204320\|ref\|NR_157735.1\| | *Bacillus proteolyticus* | firmicutes | 98.951 | 858 | 6 | 3 | 23 | 878 | 567 | 1423 | 0 | 1502 | 97 |
| 18 | gi\|1441204320\|ref\|NR_157735.1\| | *Bacillus proteolyticus* | firmicutes | 99.32 | 883 | 4 | 2 | 16 | 896 | 566 | 1448 | 0 | 1563 | 98 |
| 20 | gi\|645321690\|ref\|NR_118596.1\| | *Dietzia maris* | high G+C Gram-positive bacteria | 99.187 | 861 | 6 | 1 | 22 | 881 | 474 | 1334 | 0 | 1524 | 97 |
| 29 | gi\|659364953\|ref\|NR_121761.1\| | *Bacillus toyonensis* | firmicutes | 97.967 | 984 | 16 | 4 | 12 | 993 | 563 | 1544 | 0 | 1684 | 99 |
| 38 | gi\|310975161\|ref\|NR_037025.1\| | *Dietzia maris* | high G+C Gram-positive bacteria | 99.251 | 935 | 4 | 3 | 16 | 948 | 514 | 1447 | 0 | 1648 | 98 |
| 51 | gi\|1441204320\|ref\|NR_157735.1\| | *Bacillus proteolyticus* | firmicutes | 99.678 | 933 | 2 | 1 | 15 | 946 | 564 | 1496 | 0 | 1668 | 98 |
| 56 | gi\|636559893\|ref\|NR_115953.1\| | *Priestia aryabhattai* B8W22 | firmicutes | 98.4 | 875 | 13 | 1 | 18 | 891 | 569 | 1443 | 0 | 1523 | 98 |

**Supplemental Table S2 (continued):** Gulf of Mexico sediment bacterial isolates' BLASTn results (3/14).

| Isolate | Result_ID | Result_Sci_Names | Subject_Blast_Names | %ID | ALIGN_LEN | Mismatches | Gap Opens | Q. start | Q. end | S. start | S. end | Evalue | Bit Score | % Query Coverage Per Subject |
|---|---|---|---|---|---|---|---|---|---|---|---|---|---|---|
| 63 | gi\|566084929\|ref\|NR_108473.1\| | *Paenisporosarcina indica* | firmicutes | 96.076 | 943 | 30 | 7 | 15 | 956 | 566 | 1502 | 0 | 1528 | 98 |
| 64 | gi\|636560458\|ref\|NR_116518.1\| | *Fictibacillus rigui* | firmicutes | 99.648 | 284 | 0 | 1 | 13 | 295 | 538 | 821 | 1.23E-142 | 505 | 96 |
| 66 | gi\|636559893\|ref\|NR_115953.1\| | *Priestia aryabhattai* B8W22 | firmicutes | 98.242 | 967 | 11 | 5 | 13 | 974 | 566 | 1531 | 0 | 1661 | 93 |
| 67 | gi\|343202598\|ref\|NR_042974.1\| | *Metabacillus indicus* | firmicutes | 98.205 | 947 | 15 | 2 | 21 | 966 | 533 | 1478 | 0 | 1638 | 92 |
| 72 | gi\|645321690\|ref\|NR_118596.1\| | *Dietzia maris* | high G+C Gram-positive bacteria | 98.854 | 698 | 7 | 1 | 17 | 713 | 471 | 1168 | 0 | 1223 | 97 |
| 74 | gi\|1441204320\|ref\|NR_157735.1\| | *Bacillus proteolyticus* | firmicutes | 99.367 | 632 | 2 | 2 | 15 | 645 | 564 | 1194 | 0 | 1117 | 98 |
| 75 | gi\|636560458\|ref\|NR_116518.1\| | *Fictibacillus rigui* | firmicutes | 97.712 | 743 | 14 | 3 | 13 | 752 | 540 | 1282 | 0 | 1265 | 98 |
| 78 | gi\|636559893\|ref\|NR_115953.1\| | *Priestia aryabhattai* B8W22 | firmicutes | 98.847 | 954 | 9 | 2 | 15 | 966 | 567 | 1520 | 0 | 1672 | 90 |
| 79 | gi\|1230874627\|ref\|NR_148787.1\| | *Bacillus australimaris* | firmicutes | 98.529 | 952 | 10 | 3 | 15 | 962 | 562 | 1513 | 0 | 1651 | 88 |
| 81 | gi\|636559893\|ref\|NR_115953.1\| | *Priestia aryabhattai* B8W22 | firmicutes | 98.33 | 958 | 11 | 5 | 12 | 965 | 566 | 1522 | 0 | 1648 | 92 |
| 82 | gi\|1212229176\|ref\|NR_148244.1\| | *Bacillus xiamenensis* | firmicutes | 96.771 | 929 | 25 | 5 | 13 | 939 | 562 | 1487 | 0 | 1540 | 96 |
| 84 | gi\|1146059143\|ref\|NR_145875.1\| | *Pseudorhizobium marinum* | a-proteobacteria | 98.626 | 946 | 11 | 2 | 13 | 957 | 494 | 1438 | 0 | 1650 | 87 |

**Supplemental Table S2 (continued):** Gulf of Mexico sediment bacterial isolates' BLASTn results (4/14).

| Isolate | Result_ID | Result_Sci_Names | Subject_Blast_Names | %ID | ALIGN_LEN | Mismatches | Gap Opens | Q. start | Q. end | S. start | S. end | Evalue | Bit Score | % Query Coverage Per Subject |
|---|---|---|---|---|---|---|---|---|---|---|---|---|---|---|
| 102 | gi\|636559719\|ref\|NR_115779.1\| | *Streptomyces libani* | high G+C Gram-positive bacteria | 93.429 | 974 | 59 | 4 | 15 | 983 | 526 | 1499 | 0 | 1491 | 97 |
| 106 | gi\|631250995\|ref\|NR_112192.1\| | *Arthrobacter globiformis* | high G+C Gram-positive bacteria | 96.304 | 974 | 28 | 6 | 15 | 986 | 509 | 1476 | 0 | 1596 | 96 |
| 107 | gi\|636559893\|ref\|NR_115953.1\| | *Priestia aryabhattai* B8W22 | firmicutes | 99.373 | 957 | 4 | 2 | 14 | 968 | 566 | 1522 | 0 | 1696 | 96 |
| 108 | gi\|636559893\|ref\|NR_115953.1\| | *Priestia aryabhattai* B8W22 | firmicutes | 98.964 | 965 | 7 | 3 | 17 | 979 | 568 | 1531 | 0 | 1691 | 98 |
| 110 | gi\|636559893\|ref\|NR_115953.1\| | *Priestia aryabhattai* B8W22 | firmicutes | 98.34 | 964 | 11 | 5 | 18 | 977 | 568 | 1530 | 0 | 1659 | 98 |
| 111 | gi\|636560458\|ref\|NR_116518.1\| | *Fictibacillus rigui* | firmicutes | 97.93 | 918 | 15 | 4 | 15 | 930 | 538 | 1453 | 0 | 1570 | 92 |
| 116 | gi\|636560458\|ref\|NR_116518.1\| | *Fictibacillus rigui* | firmicutes | 99.644 | 281 | 0 | 1 | 18 | 297 | 540 | 820 | 5.30E-141 | 499 | 94 |
| 117 | gi\|1146059143\|ref\|NR_145875.1\| | *Pseudorhizobium marinum* | a-proteobacteria | 97.985 | 943 | 16 | 3 | 17 | 956 | 494 | 1436 | 0 | 1618 | 92 |
| 128 | gi\|343202598\|ref\|NR_042974.1\| | *Metabacillus indicus* | firmicutes | 96.458 | 960 | 26 | 7 | 21 | 977 | 533 | 1487 | 0 | 1576 | 85 |
| 136 | gi\|1146059154\|ref\|NR_145886.1\| | *Rhodococcus qingshengii* | high G+C Gram-positive bacteria | 82.458 | 895 | 151 | 6 | 22 | 913 | 511 | 1402 | 0 | 978 | 82 |

**Supplemental Table S2 (continued):** Gulf of Mexico sediment bacterial isolates' BLASTn results (5/14).

| Isolate | Result_ID | Result_Sci_Names | Subject_Blast_Names | %ID | ALIGN_LEN | Mismatches | Gap Opens | Q. start | Q. end | S. start | S. end | Evalue | Bit Score | % Query Coverage Per Subject |
|---|---|---|---|---|---|---|---|---|---|---|---|---|---|---|
| 142 | gi\|1315204046\|ref\|NR_152716.1\| | *Mesorhizobium sediminum* | a-proteobacteria | 98.774 | 979 | 9 | 3 | 21 | 997 | 504 | 1481 | 0 | 1709 | 85 |
| 152 | gi\|636560042\|ref\|NR_116102.1\| | *Aliihoeflea aestuarii* | a-proteobacteria | 97.679 | 948 | 21 | 1 | 18 | 964 | 496 | 1443 | 0 | 1612 | 86 |
| 169 | gi\|1315204046\|ref\|NR_152716.1\| | *Mesorhizobium sediminum* | a-proteobacteria | 98.981 | 981 | 4 | 6 | 23 | 1000 | 504 | 1481 | 0 | 1706 | 89 |
| 173 | gi\|631252086\|ref\|NR_113284.1\| | *Cellulomonas oligotrophica* | high G+C Gram-positive bacteria | 96.778 | 931 | 26 | 4 | 22 | 950 | 515 | 1443 | 0 | 1544 | 90 |
| 174 | gi\|559795186\|ref\|NR_104775.1\| | *Mycolicibacterium fortuitum* subsp. *acetamidolyticum* | high G+C Gram-positive bacteria | 97.322 | 971 | 18 | 7 | 19 | 981 | 525 | 1495 | 0 | 1624 | 84 |
| 175 | gi\|566085441\|ref\|NR_109481.1\| | *Rhodococcus nanhaiensis* | high G+C Gram-positive bacteria | 89.329 | 581 | 49 | 13 | 20 | 587 | 477 | 1057 | 0 | 754 | 97 |
| 176 | gi\|1146059143\|ref\|NR_145875.1\| | *Pseudorhizobium marinum* | a-proteobacteria | 98.726 | 942 | 6 | 5 | 22 | 962 | 500 | 1436 | 0 | 1631 | 89 |
| 177 | gi\|1146059143\|ref\|NR_145875.1\| | *Pseudorhizobium marinum* | a-proteobacteria | 99.154 | 946 | 6 | 2 | 21 | 964 | 499 | 1444 | 0 | 1667 | 88 |
| 178 | gi\|1146059143\|ref\|NR_145875.1\| | *Pseudorhizobium marinum* | a-proteobacteria | 98.946 | 949 | 6 | 4 | 21 | 968 | 499 | 1444 | 0 | 1656 | 85 |

**Supplemental Table S2 (continued):** Gulf of Mexico sediment bacterial isolates' BLASTn results (6/14).

| Isolate | Result_ID | Result_Sci_Names | Subject_Blast_Names | %ID | ALIGN_LEN | Mismatches | Gap Opens | Q. start | Q. end | S. start | S. end | Evalue | Bit Score | % Query Coverage Per Subject |
|---|---|---|---|---|---|---|---|---|---|---|---|---|---|---|
| 179 | gi\|1146059143\|ref\|NR_145875.1\| | *Pseudorhizobium marinum* | a-proteobacteria | 96.951 | 951 | 22 | 4 | 24 | 973 | 500 | 1444 | 0 | 1591 | 86 |
| 216 | gi\|559795186\|ref\|NR_104775.1\| | *Mycolicibacterium fortuitum* subsp. *acetamidolyticum* | high G+C Gram-positive bacteria | 98.68 | 985 | 12 | 1 | 15 | 998 | 521 | 1505 | 0 | 1724 | 91 |
| 219 | gi\|1146059143\|ref\|NR_145875.1\| | *Pseudorhizobium marinum* | a-proteobacteria | 81.003 | 937 | 175 | 3 | 23 | 957 | 500 | 1435 | 0 | 973 | 86 |
| 237 | gi\|219857551\|ref\|NR_025139.1\| | *Pseudoalteromonas issachenkonii* | g-proteobacteria | 95.112 | 982 | 39 | 9 | 23 | 1000 | 470 | 1446 | 0 | 1554 | 89 |
| 239 | gi\|343200197\|ref\|NR_040884.1\| | *Paenibacillus illinoisensis* | firmicutes | 97.581 | 951 | 18 | 4 | 31 | 981 | 556 | 1501 | 0 | 1612 | 83 |
| 240 | gi\|219857551\|ref\|NR_025139.1\| | *Pseudoalteromonas issachenkonii* | g-proteobacteria | 98.271 | 983 | 7 | 9 | 23 | 1001 | 470 | 1446 | 0 | 1671 | 86 |
| 250 | gi\|343201613\|ref\|NR_042339.1\| | *Bacillus aerophilus* | firmicutes | 97.339 | 977 | 14 | 8 | 22 | 995 | 564 | 1531 | 0 | 1631 | 86 |
| 4 | gi\|343202598\|ref\|NR_042974.1\| | *Metabacillus indicus* | firmicutes | 99.391 | 821 | 4 | 1 | 21 | 840 | 532 | 1352 | 0 | 1459 | 97 |
| 7 | gi\|631252797\|ref\|NR_113995.1\| | *Rossellomorea vietnamensis* | firmicutes | 99.069 | 859 | 6 | 2 | 18 | 875 | 352 | 1209 | 0 | 1512 | 98 |
| 9 | gi\|631252797\|ref\|NR_113995.1\| | *Rossellomorea vietnamensis* | firmicutes | 99.416 | 856 | 4 | 1 | 21 | 875 | 353 | 1208 | 0 | 1522 | 98 |

**Supplemental Table S2 (continued):** Gulf of Mexico sediment bacterial isolates' BLASTn results (7/14).

| Isolate | Result_ID | Result_Sci_Names | Subject_Blast_Names | %ID | ALIGN_LEN | Mismatches | Gap Opens | Q. start | Q. end | S. start | S. end | Evalue | Bit Score | % Query Coverage Per Subject |
|---|---|---|---|---|---|---|---|---|---|---|---|---|---|---|
| 25 | gi\|1240411926\|ref\|NR_149270.1\| | *Rhodococcus pedocola* | high G+C Gram-positive bacteria | 98.827 | 938 | 10 | 1 | 17 | 953 | 518 | 1455 | 0 | 1645 | 98 |
| 30 | gi\|1212229176\|ref\|NR_148244.1\| | *Bacillus xiamenensis* | firmicutes | 99.892 | 927 | 0 | 1 | 17 | 942 | 563 | 1489 | 0 | 1664 | 97 |
| 31 | gi\|1212229176\|ref\|NR_148244.1\| | *Bacillus xiamenensis* | firmicutes | 99.889 | 898 | 0 | 1 | 17 | 913 | 562 | 1459 | 0 | 1612 | 98 |
| 32 | gi\|343202598\|ref\|NR_042974.1\| | *Metabacillus indicus* | firmicutes | 99.431 | 879 | 4 | 1 | 18 | 895 | 529 | 1407 | 0 | 1563 | 98 |
| 33 | gi\|1240411945\|ref\|NR_149289.1\| | *Fictibacillus halophilus* | firmicutes | 99.144 | 935 | 5 | 3 | 22 | 954 | 513 | 1446 | 0 | 1644 | 92 |
| 35 | gi\|1441204320\|ref\|NR_157735.1\| | *Bacillus proteolyticus* | firmicutes | 99.569 | 927 | 2 | 2 | 19 | 944 | 567 | 1492 | 0 | 1649 | 98 |
| 39 | gi\|1240411945\|ref\|NR_149289.1\| | *Fictibacillus halophilus* | firmicutes | 99.351 | 925 | 5 | 1 | 13 | 936 | 511 | 1435 | 0 | 1641 | 99 |
| 41 | gi\|974142275\|ref\|NR_134795.1\| | *[Pseudomonas] zhaodongensis* | g-proteobacteria | 99.887 | 883 | 0 | 1 | 13 | 894 | 500 | 1382 | 0 | 1585 | 98 |
| 42 | gi\|974142275\|ref\|NR_134795.1\| | *[Pseudomonas] zhaodongensis* | g-proteobacteria | 99.765 | 851 | 1 | 1 | 21 | 870 | 505 | 1355 | 0 | 1524 | 98 |
| 44 | gi\|636559893\|ref\|NR_115953.1\| | *Priestia aryabhattai* B8W22 | firmicutes | 99.67 | 908 | 2 | 1 | 15 | 921 | 567 | 1474 | 0 | 1623 | 98 |
| 45 | gi\|1441204320\|ref\|NR_157735.1\| | *Bacillus proteolyticus* | firmicutes | 98.937 | 941 | 9 | 1 | 13 | 952 | 564 | 1504 | 0 | 1655 | 98 |

**Supplemental Table S2 (continued):** Gulf of Mexico sediment bacterial isolates' BLASTn results (8/14).

| Isolate | Result_ID | Result_Sci_Names | Subject_Blast_Names | %ID | ALIGN_LEN | Mismatches | Gap Opens | Q. start | Q. end | S. start | S. end | Evalue | Bit Score | % Query Coverage Per Subject |
|---|---|---|---|---|---|---|---|---|---|---|---|---|---|---|
| 46 | gi\|1441204321\|ref\|NR_157736.1\| | *Bacillus tropicus* | firmicutes | 78.925 | 930 | 187 | 7 | 23 | 948 | 575 | 1499 | 0 | 873 | 97 |
| 52 | gi\|1779814807\|ref\|NR_164882.1\| | *Bacillus zanthoxyli* | firmicutes | 98.542 | 686 | 9 | 1 | 12 | 696 | 524 | 1209 | 0 | 1195 | 72 |
| 57 | gi\|219857646\|ref\|NR_025235.1\| | *Mycolicibacterium poriferae* | high G+C Gram-positive bacteria | 93.636 | 880 | 44 | 11 | 16 | 883 | 507 | 1386 | 0 | 1333 | 87 |
| 58 | gi\|343203923\|ref\|NR_043720.1\| | *Paenisporosarcina quisquiliarum* | firmicutes | 99.664 | 893 | 2 | 1 | 14 | 905 | 551 | 1443 | 0 | 1595 | 96 |
| 59 | gi\|1315204046\|ref\|NR_152716.1\| | *Mesorhizobium sediminum* | a-proteobacteria | 99.527 | 634 | 2 | 1 | 21 | 653 | 504 | 1137 | 0 | 1128 | 97 |
| 73 | gi\|219846133\|ref\|NR_025723.1\| | *Kocuria marina* | high G+C Gram-positive bacteria | 98.889 | 540 | 5 | 1 | 14 | 552 | 519 | 1058 | 0 | 947 | 98 |
| 76 | gi\|636560641\|ref\|NR_116701.1\| | *Bacillus tianmuensis* | firmicutes | 97.959 | 980 | 15 | 4 | 12 | 986 | 562 | 1541 | 0 | 1667 | 92 |

**Supplemental Table S2 (continued):** Gulf of Mexico sediment bacterial isolates' BLASTn results (9/14).

| Isolate | Result_ID | Result_Sci_Names | Subject_Blast_Names | %ID | ALIGN_LEN | Mismatches | Gap Opens | Q. start | Q. end | S. start | S. end | Evalue | Bit Score | % Query Coverage Per Subject |
|---|---|---|---|---|---|---|---|---|---|---|---|---|---|---|
| 77 | gi\|1315204046\|ref\|NR_152716.1\| | *Mesorhizobium sediminum* | a-proteobacteria | 97.313 | 856 | 19 | 4 | 22 | 875 | 504 | 1357 | 0 | 1442 | 96 |
| 80 | gi\|1227086176\|ref\|NR_148649.1\| | *Enterobacter bugandensis* | enterobacteria | 99.028 | 926 | 7 | 2 | 17 | 940 | 512 | 1437 | 0 | 1628 | 98 |
| 83 | gi\|636559893\|ref\|NR_115953.1\| | *Priestia aryabhattai* B8W22 | firmicutes | 98.328 | 957 | 12 | 3 | 13 | 965 | 566 | 1522 | 0 | 1654 | 89 |
| 85 | gi\|636559893\|ref\|NR_115953.1\| | *Priestia aryabhattai* B8W22 | firmicutes | 98.958 | 960 | 8 | 2 | 12 | 969 | 566 | 1525 | 0 | 1687 | 90 |
| 87 | gi\|1441204321\|ref\|NR_157736.1\| | *Bacillus tropicus* | firmicutes | 96.644 | 894 | 25 | 4 | 13 | 901 | 564 | 1457 | 0 | 1485 | 97 |
| 88 | gi\|636560518\|ref\|NR_116578.1\| | *Micrococcus yunnanensis* | high G+C Gram-positive bacteria | 97.432 | 701 | 17 | 1 | 16 | 715 | 485 | 1185 | 0 | 1196 | 98 |
| 89 | gi\|1441204320\|ref\|NR_157735.1\| | *Bacillus proteolyticus* | firmicutes | 98.723 | 783 | 7 | 3 | 13 | 794 | 564 | 1344 | 0 | 1363 | 98 |
| 90 | gi\|631250995\|ref\|NR_112192.1\| | *Arthrobacter globiformis* | high G+C Gram-positive bacteria | 97.725 | 967 | 16 | 4 | 14 | 978 | 509 | 1471 | 0 | 1643 | 94 |
| 91 | gi\|636559719\|ref\|NR_115779.1\| | *Streptomyces libani* | high G+C Gram-positive bacteria | 98.973 | 974 | 7 | 3 | 16 | 986 | 525 | 1498 | 0 | 1706 | 97 |

**Supplemental Table S2 (continued):** Gulf of Mexico sediment bacterial isolates' BLASTn results (10/14).

| Isolate | Result_ID | Result_Sci_Names | Subject_Blast_Names | %ID | ALIGN_LEN | Mismatches | Gap Opens | Q. start | Q. end | S. start | S. end | Evalue | Bit Score | % Query Coverage Per Subject |
|---|---|---|---|---|---|---|---|---|---|---|---|---|---|---|
| 93 | gi\|219857652\|ref\|NR_025241.1\| | *Rossellomorea aquimaris* | firmicutes | 98.233 | 962 | 13 | 3 | 16 | 975 | 543 | 1502 | 0 | 1658 | 94 |
| 94 | gi\|343201613\|ref\|NR_042339.1\| | *Bacillus aerophilus* | firmicutes | 98.765 | 972 | 9 | 2 | 16 | 984 | 560 | 1531 | 0 | 1700 | 94 |
| 96 | gi\|636559893\|ref\|NR_115953.1\| | *Priestia aryabhattai* B8W22 | firmicutes | 97.619 | 966 | 20 | 3 | 15 | 977 | 566 | 1531 | 0 | 1645 | 91 |
| 97 | gi\|1269807446\|ref\|NR_074714.2\| | *Kytococcus sedentarius* | high G+C Gram-positive bacteria | 97.459 | 984 | 17 | 7 | 19 | 996 | 543 | 1524 | 0 | 1653 | 97 |
| 98 | gi\|636559893\|ref\|NR_115953.1\| | *Priestia aryabhattai* B8W22 | firmicutes | 99.069 | 967 | 7 | 2 | 13 | 978 | 566 | 1531 | 0 | 1703 | 96 |
| 101 | gi\|343201613\|ref\|NR_042339.1\| | *Bacillus aerophilus* | firmicutes | 99.486 | 972 | 3 | 2 | 15 | 984 | 560 | 1531 | 0 | 1727 | 98 |
| 105 | gi\|219857651\|ref\|NR_025240.1\| | *Rossellomorea marisflavi* | firmicutes | 99.582 | 958 | 3 | 1 | 19 | 975 | 544 | 1501 | 0 | 1709 | 91 |
| 113 | gi\|343201613\|ref\|NR_042339.1\| | *Bacillus aerophilus* | firmicutes | 98.664 | 973 | 8 | 5 | 16 | 984 | 560 | 1531 | 0 | 1686 | 94 |
| 114 | gi\|659364953\|ref\|NR_121761.1\| | *Bacillus toyonensis* | firmicutes | 98.883 | 985 | 8 | 3 | 15 | 997 | 561 | 1544 | 0 | 1723 | 96 |
| 115 | gi\|636559893\|ref\|NR_115953.1\| | *Priestia aryabhattai* B8W22 | firmicutes | 98.858 | 963 | 7 | 4 | 16 | 974 | 569 | 1531 | 0 | 1679 | 91 |

**Supplemental Table S2 (continued):** Gulf of Mexico sediment bacterial isolates' BLASTn results (11/14).

| Isolate | Result_ID | Result_Sci_Names | Subject_Blast_Names | %ID | ALIGN_LEN | Mismatches | Gap Opens | Q. start | Q. end | S. start | S. end | Evalue | Bit Score | % Query Coverage Per Subject |
|---|---|---|---|---|---|---|---|---|---|---|---|---|---|---|
| 118 | gi\|310975161\|ref\|NR_037025.1\| | *Dietzia maris* | high G+C Gram-positive bacteria | 98.647 | 961 | 9 | 3 | 18 | 976 | 515 | 1473 | 0 | 1673 | 98 |
| 121 | gi\|1146059143\|ref\|NR_145875.1\| | *Pseudorhizobium marinum* | a-proteobacteria | 85.448 | 536 | 70 | 7 | 16 | 545 | 498 | 1031 | 1.94E-178 | 626 | 52 |
| 123 | gi\|1240411945\|ref\|NR_149289.1\| | *Fictibacillus halophilus* | firmicutes | 96.133 | 905 | 27 | 4 | 61 | 964 | 549 | 1446 | 0 | 1487 | 86 |
| 130 | gi\|1146059154\|ref\|NR_145886.1\| | *Rhodococcus qingshengii* | high G+C Gram-positive bacteria | 79.218 | 895 | 166 | 16 | 20 | 905 | 511 | 1394 | 0 | 797 | 80 |
| 133 | gi\|1315204046\|ref\|NR_152716.1\| | *Mesorhizobium sediminum* | a-proteobacteria | 97.541 | 976 | 20 | 4 | 21 | 994 | 504 | 1477 | 0 | 1654 | 95 |
| 134 | gi\|1315204046\|ref\|NR_152716.1\| | *Mesorhizobium sediminum* | a-proteobacteria | 96.957 | 986 | 25 | 5 | 16 | 1000 | 499 | 1480 | 0 | 1643 | 95 |
| 135 | gi\|1146059154\|ref\|NR_145886.1\| | *Rhodococcus qingshengii* | high G+C Gram-positive bacteria | 71.991 | 432 | 112 | 6 | 72 | 498 | 562 | 989 | 2.51E-63 | 243 | 40 |
| 137 | gi\|1315204046\|ref\|NR_152716.1\| | *Mesorhizobium sediminum* | a-proteobacteria | 98.679 | 984 | 9 | 4 | 16 | 996 | 499 | 1481 | 0 | 1708 | 94 |

**Supplemental Table S2 (continued):** Gulf of Mexico sediment bacterial isolates' BLASTn results (12/14).

| Isolate | Result_ID | Result_Sci_Names | Subject_Blast_Names | %ID | ALIGN_LEN | Mismatches | Gap Opens | Q. start | Q. end | S. start | S. end | Evalue | Bit Score | % Query Coverage Per Subject |
|---|---|---|---|---|---|---|---|---|---|---|---|---|---|---|
| 140 | gi\|636559761\|ref\|NR_115821.1\| | *Alcanivorax pacificus* W11-5 | g-proteobacteria | 92.479 | 944 | 62 | 9 | 21 | 961 | 559 | 1496 | 0 | 1379 | 88 |
| 143 | gi\|559795186\|ref\|NR_104775.1\| | *Mycolicibacterium fortuitum* subsp. *acetamidolyticum* | high G+C Gram-positive bacteria | 98.68 | 985 | 11 | 2 | 15 | 997 | 521 | 1505 | 0 | 1720 | 86 |
| 146 | gi\|559795186\|ref\|NR_104775.1\| | *Mycolicibacterium fortuitum* subsp. *acetamidolyticum* | high G+C Gram-positive bacteria | 98.777 | 981 | 9 | 3 | 24 | 1002 | 526 | 1505 | 0 | 1712 | 90 |
| 147 | gi\|559795186\|ref\|NR_104775.1\| | *Mycolicibacterium fortuitum* subsp. *acetamidolyticum* | high G+C Gram-positive bacteria | 98.473 | 982 | 10 | 5 | 18 | 994 | 524 | 1505 | 0 | 1694 | 90 |
| 148 | gi\|1315204046\|ref\|NR_152716.1\| | *Mesorhizobium sediminum* | a-proteobacteria | 98.978 | 978 | 8 | 2 | 19 | 994 | 504 | 1481 | 0 | 1718 | 89 |
| 153 | gi\|1491514589\|ref\|NR_159197.1\| | *Mycobacterium neumannii* | high G+C Gram-positive bacteria | 85.228 | 941 | 126 | 10 | 19 | 952 | 541 | 1475 | 0 | 1100 | 87 |
| 156 | gi\|1315204046\|ref\|NR_152716.1\| | *Mesorhizobium sediminum* | a-proteobacteria | 98.632 | 950 | 10 | 3 | 22 | 968 | 504 | 1453 | 0 | 1651 | 91 |

**Supplemental Table S2 (continued):** Gulf of Mexico sediment bacterial isolates' BLASTn results (13/14).

| Isolate | Result_ID | Result_Sci_Names | Subject_Blast_Names | %ID | ALIGN_LEN | Mismatches | Gap Opens | Q. start | Q. end | S. start | S. end | Evalue | Bit Score | % Query Coverage Per Subject |
|---|---|---|---|---|---|---|---|---|---|---|---|---|---|---|
| 162 | gi\|559795186\|ref\|NR_104775.1\| | *Mycolicibacterium fortuitum* subsp. *acetamidolyticum* | high G+C Gram-positive bacteria | 97.439 | 976 | 16 | 5 | 16 | 982 | 522 | 1497 | 0 | 1647 | 88 |
| 163 | gi\|559795186\|ref\|NR_104775.1\| | *Mycolicibacterium fortuitum* subsp. *acetamidolyticum* | high G+C Gram-positive bacteria | 97.546 | 978 | 17 | 6 | 19 | 991 | 524 | 1499 | 0 | 1649 | 91 |
| 164 | gi\|559795186\|ref\|NR_104775.1\| | *Mycolicibacterium fortuitum* subsp. *acetamidolyticum* | high G+C Gram-positive bacteria | 98.463 | 976 | 13 | 2 | 18 | 991 | 522 | 1497 | 0 | 1697 | 91 |
| 165 | gi\|559795186\|ref\|NR_104775.1\| | *Mycolicibacterium fortuitum* subsp. *acetamidolyticum* | high G+C Gram-positive bacteria | 98.663 | 972 | 10 | 3 | 25 | 993 | 526 | 1497 | 0 | 1692 | 90 |
| 166 | gi\|1315204046\|ref\|NR_152716.1\| | *Mesorhizobium sediminum* | a-proteobacteria | 99.08 | 978 | 7 | 2 | 21 | 996 | 504 | 1481 | 0 | 1722 | 90 |
| 167 | gi\|559795186\|ref\|NR_104775.1\| | *Mycolicibacterium fortuitum* subsp. *acetamidolyticum* | high G+C Gram-positive bacteria | 94.731 | 968 | 48 | 3 | 35 | 999 | 538 | 1505 | 0 | 1527 | 91 |
| 168 | gi\|636559761\|ref\|NR_115821.1\| | *Alcanivorax pacificus* W11-5 | g-proteobacteria | 97.808 | 958 | 6 | 15 | 23 | 974 | 558 | 1506 | 0 | 1585 | 80 |

**Supplemental Table S2 (continued):** Gulf of Mexico sediment bacterial isolates' BLASTn results (14/14).

| Isolate | Result_ID | Result_Sci_Names | Subject_Blast_Names | %ID | ALIGN_LEN | Mismatches | Gap Opens | Q. start | Q. end | S. start | S. end | Evalue | Bit Score | % Query Coverage Per Subject |
|---|---|---|---|---|---|---|---|---|---|---|---|---|---|---|
| 217 | gi\|559795186\|ref\|NR_104775.1\| | *Mycolicibacterium fortuitum* subsp. *acetamidolyticum* | high G+C Gram-positive bacteria | 98.777 | 981 | 9 | 3 | 21 | 999 | 526 | 1505 | 0 | 1712 | 83 |
| 218 | gi\|559795186\|ref\|NR_104775.1\| | *Mycolicibacterium fortuitum* subsp. *acetamidolyticum* | high G+C Gram-positive bacteria | 98.984 | 984 | 7 | 3 | 17 | 997 | 522 | 1505 | 0 | 1725 | 86 |

**Supplemental Table S3.** Recovered Antarctic sediment-associated taxonomy spans 3 phyla.

| Genera | Family | Order | Class | Phylum |
|---|---|---|---|---|
| *Agrococcus* | *Microbacteriaceae* | Micrococcales | Actinomycetes | Actinomycetota |
| *Alkalihalobacillus* | *Bacillaceae* | Caryophanales | Bacilli | Bacillota |
| *Bacillus* | *Bacillaceae* | Caryophanales | Bacilli | Bacillota |
| *Bhargavaea* | *Caryophanaceae* | Caryophanales | Bacilli | Bacillota |
| *Cytobacillus* | *Bacillaceae* | Caryophanales | Bacilli | Bacillota |
| *Dietzia* | *Dietziaceae* | Mycobacteriales | Actinomycetes | Actinomycetota |
| *Fictibacillus* | *Bacillaceae* | Caryophanales | Bacilli | Bacillota |
| *Mesobacillus* | *Bacillaceae* | Caryophanales | Bacilli | Bacillota |
| *Metabacillus* | *Bacillaceae* | Caryophanales | Bacilli | Bacillota |
| *Micrococcus* | *Micrococcaceae* | Micrococcales | Actinomycetes | Actinomycetota |
| *Nocardioides* | *Nocardioidaceae* | Propionibacteriales | Actinomycetes | Actinomycetota |
| *Ornithinicoccus* | *Intrasporangiaceae* | Micrococcales | Actinomycetes | Actinomycetota |
| *Ornithinimicrobium* | *Ornithinimicrobiaceae* | Micrococcales | Actinomycetes | Actinomycetota |
| *Paenisporosarcina* | *Caryophanaceae* | Caryophanales | Bacilli | Bacillota |
| *Psychrobacter* | *Moraxellaceae* | Pseudomonadales | Gammaproteobacteria | Pseudomonadota |
| *Rhodococcus* | *Nocardiaceae* | Mycobacteriales | Actinomycetes | Actinomycetota |
| *Robertmurraya* | *Bacillaceae* | Caryophanales | Bacilli | Bacillota |
| *Rossellomorea* | *Bacillaceae* | Caryophanales | Bacilli | Bacillota |
| *Sporosarcina* | *Caryophanaceae* | Caryophanales | Bacilli | Bacillota |
| *Tomitella* | *Nocardiaceae* | Mycobacteriales | Actinomycetes | Actinomycetota |
| *Virgibacillus* | *Bacillaceae* | Caryophanales | Bacilli | Bacillota |

**Supplemental Table S4.** Antarctic bacteria from 20-ft (TT-abbreviated) and 60-ft BLASTn results (1/17).

| Isolate | Result_ID | Result_Sci_Names | Subject_Blast_Names | %ID | ALIGN_LEN | Mismatches | Gap Opens | Q. start | Q. end | S. start | S. end | Evalue | Bit Score | % Query Coverage Per Subject |
|---|---|---|---|---|---|---|---|---|---|---|---|---|---|---|
| TT-144 | gi\|631251707\|ref\|NR_112905.1\| | *Tomitella biformata* AHU 1821 | high G+C Gram-positive bacteria | 96.381 | 967 | 30 | 5 | 18 | 982 | 508 | 1471 | 0 | 1583 | 87 |
| TT-145 | gi\|444439739\|ref\|NR_075054.1\| | *Psychrobacter arcticus* | g-proteobacteria | 96.751 | 985 | 21 | 9 | 19 | 1002 | 608 | 1582 | 0 | 1610 | 90 |
| TT-146 | gi\|636560789\|ref\|NR_116849.1\| | *Robertmurraya beringensis* | firmicutes | 91.336 | 554 | 44 | 4 | 19 | 568 | 564 | 1117 | 0 | 783 | 50 |
| TT-147 | gi\|645319993\|ref\|NR_117306.1\| | *Dietzia alimentaria* 72 | high G+C Gram-positive bacteria | 98.938 | 942 | 7 | 3 | 21 | 960 | 484 | 1424 | 0 | 1647 | 85 |
| TT-148 | gi\|343202719\|ref\|NR_043141.1\| | *Psychrobacter namhaensis* | g-proteobacteria | 93.035 | 962 | 64 | 3 | 17 | 976 | 534 | 1494 | 0 | 1455 | 85 |
| TT-24 | gi\|444439739\|ref\|NR_075054.1\| | *Psychrobacter arcticus* | g-proteobacteria | 96.862 | 988 | 26 | 4 | 15 | 1001 | 602 | 1585 | 0 | 1642 | 89 |
| TT-35 | gi\|343201095\|ref\|NR_041782.1\| | *Sporosarcina ureae* | firmicutes | 99.073 | 971 | 4 | 4 | 14 | 983 | 541 | 1507 | 0 | 1700 | 86 |
| TT-38 | gi\|343201095\|ref\|NR_041782.1\| | *Sporosarcina ureae* | firmicutes | 99.378 | 965 | 5 | 1 | 19 | 982 | 543 | 1507 | 0 | 1715 | 90 |
| TT-39 | gi\|645319993\|ref\|NR_117306.1\| | *Dietzia alimentaria* 72 | high G+C Gram-positive bacteria | 98.197 | 943 | 16 | 1 | 19 | 960 | 480 | 1422 | 0 | 1634 | 89 |
| TT-40 | gi\|343201095\|ref\|NR_041782.1\| | *Sporosarcina ureae* | firmicutes | 95.568 | 925 | 22 | 12 | 64 | 978 | 585 | 1500 | 0 | 1466 | 78 |
| TT-50 | gi\|636560215\|ref\|NR_116275.1\| | *Rhodococcus cercidiphylli* | high G+C Gram-positive bacteria | 98.043 | 511 | 7 | 3 | 24 | 533 | 499 | 1007 | 0 | 869 | 92 |

**Supplemental Table S4 (continued).** Antarctic bacteria from 20-ft (TT-abbreviated) and 60-ft BLASTn results (2/17).

| Isolate | Result_ID | Result_Sci_Names | Subject_Blast_Names | %ID | ALIGN_LEN | Mismatches | Gap Opens | Q. start | Q. end | S. start | S. end | Evalue | Bit Score | % Query Coverage Per Subject |
|---|---|---|---|---|---|---|---|---|---|---|---|---|---|---|
| TT-52 | gi\|343201095\|ref\|NR_041782.1\| | *Sporosarcina ureae* | firmicutes | 98.45 | 968 | 12 | 3 | 19 | 984 | 541 | 1507 | 0 | 1679 | 86 |
| TT-74 | gi\|636560215\|ref\|NR_116275.1\| | *Rhodococcus cercidiphylli* | high G+C Gram-positive bacteria | 94.513 | 966 | 43 | 7 | 40 | 1003 | 513 | 1470 | 0 | 1513 | 82 |
| TT-76 | gi\|631252554\|ref\|NR_113752.1\| | *Sporosarcina psychrophila* | firmicutes | 96.994 | 499 | 15 | 0 | 35 | 533 | 559 | 1057 | 0 | 844 | 91 |
| TT-81 | gi\|343200672\|ref\|NR_041359.1\| | *Sporosarcina saromensis* | firmicutes | 96.366 | 963 | 26 | 8 | 24 | 982 | 545 | 1502 | 0 | 1570 | 86 |
| TT-87 | gi\|343200672\|ref\|NR_041359.1\| | *Sporosarcina saromensis* | firmicutes | 98.227 | 959 | 14 | 3 | 20 | 976 | 545 | 1502 | 0 | 1653 | 89 |
| TT-88 | gi\|219857652\|ref\|NR_025241.1\| | *Rossellomorea aquimaris* | firmicutes | 99.062 | 959 | 4 | 5 | 19 | 974 | 543 | 1499 | 0 | 1675 | 88 |
| TT-89 | gi\|444439739\|ref\|NR_075054.1\| | *Psychrobacter arcticus* | g-proteobacteria | 97.862 | 982 | 15 | 6 | 21 | 1000 | 608 | 1585 | 0 | 1660 | 89 |
| TT-91 | gi\|631252554\|ref\|NR_113752.1\| | *Sporosarcina psychrophila* | firmicutes | 88.15 | 692 | 66 | 15 | 23 | 700 | 547 | 1236 | 0 | 868 | 66 |
| TT-98 | gi\|631253085\|ref\|NR_114283.1\| | *Sporosarcina luteola* | firmicutes | 80.944 | 551 | 99 | 6 | 128 | 673 | 654 | 1203 | 3.53E-150 | 532 | 50 |
| TT-99 | gi\|631253051\|ref\|NR_114249.1\| | *Sporosarcina saromensis* | firmicutes | 89.65 | 628 | 55 | 8 | 102 | 721 | 628 | 1253 | 0 | 832 | 57 |

**Supplemental Table S4 (continued).** Antarctic bacteria from 20-ft (TT-abbreviated) and 60-ft BLASTn results (3/17).

| Isolate | Result_ID | Result_Sci_Names | Subject_Blast_Names | %ID | ALIGN_LEN | Mismatches | Gap Opens | Q. start | Q. end | S. start | S. end | Evalue | Bit Score | % Query Coverage Per Subject |
|---|---|---|---|---|---|---|---|---|---|---|---|---|---|---|
| TT-101 | gi\|343201495\|ref\|NR_042221.1\| | *Psychrobacter urativorans* | g-proteobacteria | 94.838 | 988 | 36 | 11 | 23 | 1005 | 552 | 1529 | 0 | 1536 | 88 |
| TT-102 | gi\|444439739\|ref\|NR_075054.1\| | *Psychrobacter arcticus* | g-proteobacteria | 97.955 | 978 | 14 | 6 | 22 | 996 | 608 | 1582 | 0 | 1656 | 89 |
| TT-79 | gi\|444439739\|ref\|NR_075054.1\| | *Psychrobacter arcticus* | g-proteobacteria | 97.769 | 986 | 14 | 8 | 18 | 1001 | 606 | 1585 | 0 | 1661 | 85 |
| TT-149 | gi\|645319993\|ref\|NR_117306.1\| | *Dietzia alimentaria* 72 | high G+C Gram-positive bacteria | 97.576 | 949 | 15 | 7 | 22 | 966 | 485 | 1429 | 0 | 1593 | 85 |
| TT-150 | gi\|631251707\|ref\|NR_112905.1\| | *Tomitella biformata* AHU 1821 | high G+C Gram-positive bacteria | 97.704 | 958 | 19 | 3 | 23 | 977 | 511 | 1468 | 0 | 1623 | 85 |
| TT-151 | gi\|219857652\|ref\|NR_025241.1\| | *Rossellomorea aquimaris* | firmicutes | 96.751 | 954 | 23 | 6 | 33 | 985 | 556 | 1502 | 0 | 1582 | 84 |
| TT-152 | gi\|636560215\|ref\|NR_116275.1\| | *Rhodococcus cercidiphylli* | high G+C Gram-positive bacteria | 97.325 | 972 | 19 | 6 | 31 | 997 | 508 | 1477 | 0 | 1627 | 87 |
| TT-153 | gi\|636558971\|ref\|NR_115028.1\| | *Agrococcus jenensis* | high G+C Gram-positive bacteria | 97.505 | 962 | 17 | 7 | 14 | 969 | 511 | 1471 | 0 | 1612 | 86 |
| TT-154 | gi\|219846891\|ref\|NR_026483.1\| | *Ornithinicoccus hortensis* | high G+C Gram-positive bacteria | 96.923 | 975 | 23 | 6 | 17 | 984 | 518 | 1492 | 0 | 1614 | 89 |

**Supplemental Table S4 (continued).** Antarctic bacteria from 20-ft (TT-abbreviated) and 60-ft BLASTn results (4/17).

| Isolate | Result_ID | Result_Sci_Names | Subject_Blast_Names | %ID | ALIGN_LEN | Mismatches | Gap Opens | Q. start | Q. end | S. start | S. end | Evalue | Bit Score | % Query Coverage Per Subject |
|---|---|---|---|---|---|---|---|---|---|---|---|---|---|---|
| TT-155 | gi\|444439739\|ref\|NR_075054.1\| | *Psychrobacter arcticus* | g-proteobacteria | 96.057 | 989 | 23 | 13 | 22 | 1004 | 607 | 1585 | 0 | 1581 | 89 |
| TT-157 | gi\|343198706\|ref\|NR_043334.1\| | *Metabacillus niabensis* | firmicutes | 97.545 | 937 | 18 | 4 | 20 | 951 | 540 | 1476 | 0 | 1585 | 89 |
| TT-158 | gi\|636560215\|ref\|NR_116275.1\| | *Rhodococcus cercidiphylli* | high G+C Gram-positive bacteria | 98.059 | 979 | 15 | 4 | 24 | 998 | 499 | 1477 | 0 | 1674 | 87 |
| TT-159 | gi\|651343563\|ref\|NR_075062.2\| | *Micrococcus luteus* | high G+C Gram-positive bacteria | 98.784 | 987 | 9 | 3 | 19 | 1002 | 539 | 1525 | 0 | 1720 | 88 |
| TT-160 | gi\|636558971\|ref\|NR_115028.1\| | *Agrococcus jenensis* | high G+C Gram-positive bacteria | 95.248 | 505 | 24 | 0 | 25 | 529 | 524 | 1028 | 0 | 811 | 93 |
| TT-161 | gi\|265678925\|ref\|NR_029233.1\| | *Sporosarcina globispora* | firmicutes | 97.462 | 985 | 19 | 4 | 14 | 992 | 549 | 1533 | 0 | 1664 | 87 |
| TT-162 | gi\|343201095\|ref\|NR_041782.1\| | *Sporosarcina ureae* | firmicutes | 98.125 | 960 | 13 | 5 | 14 | 968 | 543 | 1502 | 0 | 1641 | 86 |
| TT-163 | gi\|343198706\|ref\|NR_043334.1\| | *Metabacillus niabensis* | firmicutes | 98.404 | 940 | 11 | 3 | 16 | 951 | 539 | 1478 | 0 | 1623 | 84 |
| TT-164 | gi\|636560215\|ref\|NR_116275.1\| | *Rhodococcus cercidiphylli* | high G+C Gram-positive bacteria | 96.939 | 980 | 24 | 5 | 20 | 993 | 498 | 1477 | 0 | 1631 | 89 |
| TT-165 | gi\|219846891\|ref\|NR_026483.1\| | *Ornithinicoccus hortensis* | high G+C Gram-positive bacteria | 97.231 | 975 | 19 | 7 | 22 | 989 | 519 | 1492 | 0 | 1621 | 91 |

**Supplemental Table S4 (continued).** Antarctic bacteria from 20-ft (TT-abbreviated) and 60-ft BLASTn results (5/17).

| Isolate | Result_ID | Result_Sci_Names | Subject_Blast_Names | %ID | ALIGN_LEN | Mismatches | Gap Opens | Q. start | Q. end | S. start | S. end | Evalue | Bit Score | % Query Coverage Per Subject |
|---|---|---|---|---|---|---|---|---|---|---|---|---|---|---|
| TT-166 | gi\|631251707\|ref\|NR_112905.1\| | *Tomitella biformata* AHU 1821 | high G+C Gram-positive bacteria | 97.083 | 960 | 23 | 4 | 18 | 972 | 509 | 1468 | 0 | 1599 | 87 |
| TT-34 | gi\|636560215\|ref\|NR_116275.1\| | *Rhodococcus cercidiphylli* | high G+C Gram-positive bacteria | 95.921 | 858 | 29 | 6 | 25 | 878 | 499 | 1354 | 0 | 1390 | 97 |
| TT-60 | gi\|631253085\|ref\|NR_114283.1\| | *Sporosarcina luteola* | firmicutes | 99.341 | 910 | 5 | 1 | 15 | 923 | 541 | 1450 | 0 | 1616 | 98 |
| TT-62 | gi\|631251646\|ref\|NR_112844.1\| | *Sporosarcina luteola* | firmicutes | 97.983 | 942 | 17 | 2 | 17 | 956 | 541 | 1482 | 0 | 1621 | 98 |
| TT-63 | gi\|343201095\|ref\|NR_041782.1\| | *Sporosarcina ureae* | firmicutes | 97.968 | 886 | 14 | 4 | 19 | 900 | 543 | 1428 | 0 | 1511 | 98 |
| TT-65 | gi\|636560215\|ref\|NR_116275.1\| | *Rhodococcus cercidiphylli* | high G+C Gram-positive bacteria | 98.169 | 874 | 14 | 2 | 17 | 889 | 495 | 1367 | 0 | 1506 | 98 |
| TT-66 | gi\|645319993\|ref\|NR_117306.1\| | *Dietzia alimentaria* 72 | high G+C Gram-positive bacteria | 99.651 | 860 | 2 | 1 | 19 | 877 | 483 | 1342 | 0 | 1535 | 98 |
| TT-67 | gi\|636560215\|ref\|NR_116275.1\| | *Rhodococcus cercidiphylli* | high G+C Gram-positive bacteria | 99.173 | 846 | 6 | 1 | 19 | 863 | 498 | 1343 | 0 | 1492 | 98 |
| TT-68 | gi\|636560518\|ref\|NR_116578.1\| | *Micrococcus yunnanensis* | high G+C Gram-positive bacteria | 99.02 | 816 | 6 | 2 | 16 | 830 | 485 | 1299 | 0 | 1434 | 98 |
| TT-70 | gi\|636560215\|ref\|NR_116275.1\| | *Rhodococcus cercidiphylli* | high G+C Gram-positive bacteria | 99.298 | 855 | 5 | 1 | 17 | 870 | 495 | 1349 | 0 | 1512 | 98 |

**Supplemental Table S4 (continued).** Antarctic bacteria from 20-ft (TT-abbreviated) and 60-ft BLASTn results (6/17).

| Isolate | Result_ID | Result_Sci_Names | Subject_Blast_Names | %ID | ALIGN_LEN | Mismatches | Gap Opens | Q. start | Q. end | S. start | S. end | Evalue | Bit Score | % Query Coverage Per Subject |
|---|---|---|---|---|---|---|---|---|---|---|---|---|---|---|
| TT-94 | gi\|636558971\|ref\|NR_115028.1\| | *Agrococcus jenensis* | high G+C Gram-positive bacteria | 98.989 | 890 | 7 | 2 | 17 | 905 | 515 | 1403 | 0 | 1561 | 98 |
| TT-61 | gi\|1240411856\|ref\|NR_149187.1\| | *Nocardioides gilvus* | high G+C Gram-positive bacteria | 97.801 | 864 | 16 | 3 | 20 | 882 | 528 | 1389 | 0 | 1472 | 98 |
| TT-111 | gi\|631252554\|ref\|NR_113752.1\| | *Sporosarcina psychrophila* | firmicutes | 95.537 | 941 | 35 | 5 | 20 | 956 | 540 | 1477 | 0 | 1519 | 96 |
| TT-138 | gi\|219857652\|ref\|NR_025241.1\| | *Rossellomorea aquimaris* | firmicutes | 98.962 | 963 | 6 | 4 | 19 | 978 | 542 | 1503 | 0 | 1683 | 95 |
| TT-11 | gi\|559795330\|ref\|NR_104923.1\| | *Sporosarcina pasteurii* | firmicutes | 97.026 | 975 | 26 | 3 | 18 | 989 | 502 | 1476 | 0 | 1630 | 98 |
| TT-90 | gi\|631251707\|ref\|NR_112905.1\| | *Tomitella biformata* AHU 1821 | high G+C Gram-positive bacteria | 96.754 | 955 | 26 | 4 | 21 | 970 | 510 | 1464 | 0 | 1581 | 98 |
| TT-69 | gi\|645319993\|ref\|NR_117306.1\| | *Dietzia alimentaria* 72 | high G+C Gram-positive bacteria | 98.079 | 937 | 15 | 3 | 21 | 955 | 484 | 1419 | 0 | 1607 | 98 |
| TT-10 | gi\|645319993\|ref\|NR_117306.1\| | *Dietzia alimentaria* 72 | high G+C Gram-positive bacteria | 98.368 | 919 | 13 | 2 | 21 | 938 | 483 | 1400 | 0 | 1590 | 98 |
| TT-15 | gi\|645319993\|ref\|NR_117306.1\| | *Dietzia alimentaria* 72 | high G+C Gram-positive bacteria | 98.089 | 942 | 16 | 2 | 21 | 961 | 485 | 1425 | 0 | 1619 | 97 |

**Supplemental Table S4 (continued).** Antarctic bacteria from 20-ft (TT-abbreviated) and 60-ft BLASTn results (7/17).

| Isolate | Result_ID | Result_Sci_Names | Subject_Blast_Names | %ID | ALIGN_LEN | Mismatches | Gap Opens | Q. start | Q. end | S. start | S. end | Evalue | Bit Score | % Query Coverage Per Subject |
|---|---|---|---|---|---|---|---|---|---|---|---|---|---|---|
| TT-84 | gi\|265678925\|ref\|NR_029233.1\| | *Sporosarcina globispora* | firmicutes | 98.319 | 952 | 13 | 3 | 16 | 966 | 549 | 1498 | 0 | 1639 | 98 |
| TT-86 | gi\|645319993\|ref\|NR_117306.1\| | *Dietzia alimentaria* 72 | high G+C Gram-positive bacteria | 94.708 | 926 | 45 | 4 | 24 | 946 | 487 | 1411 | 0 | 1471 | 94 |
| TT-100 | gi\|631251707\|ref\|NR_112905.1\| | *Tomitella biformata* AHU 1821 | high G+C Gram-positive bacteria | 94.781 | 958 | 43 | 6 | 22 | 973 | 511 | 1467 | 0 | 1508 | 90 |
| TT-22 | gi\|343205966\|ref\|NR_044482.1\| | *Dietzia kunjamensis* subsp*. schimae* | high G+C Gram-positive bacteria | 92.089 | 986 | 66 | 8 | 20 | 1002 | 535 | 1511 | 0 | 1449 | 92 |
| TT-1 | gi\|1269801501\|ref\|NR_075005.2\| | *Bacillus velezensis* | firmicutes | 98.613 | 937 | 6 | 5 | 17 | 946 | 571 | 1507 | 0 | 1619 | 87 |
| TT-4 | gi\|310975161\|ref\|NR_037025.1\| | *Dietzia maris* | high G+C Gram-positive bacteria | 97.002 | 934 | 23 | 4 | 17 | 948 | 516 | 1446 | 0 | 1564 | 87 |
| TT-7 | gi\|343202628\|ref\|NR_043009.1\| | *Rhodococcus yunnanensis* | high G+C Gram-positive bacteria | 97.983 | 942 | 16 | 2 | 16 | 956 | 506 | 1445 | 0 | 1617 | 87 |
| TT-12 | gi\|636560215\|ref\|NR_116275.1\| | *Rhodococcus cercidiphylli* | high G+C Gram-positive bacteria | 96.22 | 582 | 18 | 2 | 18 | 598 | 498 | 1076 | 0 | 957 | 97 |
| TT-17 | gi\|645321690\|ref\|NR_118596.1\| | *Dietzia maris* | high G+C Gram-positive bacteria | 99.482 | 579 | 2 | 1 | 19 | 596 | 473 | 1051 | 0 | 1029 | 97 |
| TT-33 | gi\|636560215\|ref\|NR_116275.1\| | *Rhodococcus cercidiphylli* | high G+C Gram-positive bacteria | 96.73 | 581 | 15 | 2 | 19 | 598 | 498 | 1075 | 0 | 966 | 97 |

**Supplemental Table S4 (continued).** Antarctic bacteria from 20-ft (TT-abbreviated) and 60-ft BLASTn results (8/17).

| Isolate | Result_ID | Result_Sci_Names | Subject_Blast_Names | %ID | ALIGN_LEN | Mismatches | Gap Opens | Q. start | Q. end | S. start | S. end | Evalue | Bit Score | % Query Coverage Per Subject |
|---|---|---|---|---|---|---|---|---|---|---|---|---|---|---|
| TT-64 | gi\|636560215\|ref\|NR_116275.1\| | *Rhodococcus cercidiphylli* | high G+C Gram-positive bacteria | 97.246 | 581 | 11 | 3 | 19 | 596 | 497 | 1075 | 0 | 971 | 97 |
| TT-72 | gi\|636560215\|ref\|NR_116275.1\| | *Rhodococcus cercidiphylli* | high G+C Gram-positive bacteria | 96.564 | 582 | 17 | 2 | 16 | 596 | 497 | 1076 | 0 | 965 | 97 |
| TT-75 | gi\|343202628\|ref\|NR_043009.1\| | *Rhodococcus yunnanensis* | high G+C Gram-positive bacteria | 95.69 | 928 | 32 | 6 | 16 | 938 | 506 | 1430 | 0 | 1497 | 87 |
| TT-85 | gi\|343202628\|ref\|NR_043009.1\| | *Rhodococcus yunnanensis* | high G+C Gram-positive bacteria | 94.181 | 928 | 49 | 5 | 16 | 940 | 506 | 1431 | 0 | 1451 | 86 |
| TT-103 | gi\|343200672\|ref\|NR_041359.1\| | *Sporosarcina saromensis* | firmicutes | 99.045 | 942 | 6 | 3 | 13 | 952 | 542 | 1482 | 0 | 1651 | 87 |
| TT-105 | gi\|636558971\|ref\|NR_115028.1\| | *Agrococcus jenensis* | high G+C Gram-positive bacteria | 97.497 | 959 | 22 | 2 | 15 | 971 | 515 | 1473 | 0 | 1629 | 87 |
| TT-113 | gi\|636558971\|ref\|NR_115028.1\| | *Agrococcus jenensis* | high G+C Gram-positive bacteria | 95.298 | 957 | 34 | 9 | 18 | 967 | 517 | 1469 | 0 | 1516 | 90 |
| TT-116 | gi\|631252554\|ref\|NR_113752.1\| | *Sporosarcina psychrophila* | firmicutes | 98.278 | 929 | 11 | 5 | 16 | 939 | 543 | 1471 | 0 | 1592 | 89 |
| TT-117 | gi\|636560215\|ref\|NR_116275.1\| | *Rhodococcus cercidiphylli* | high G+C Gram-positive bacteria | 96.379 | 580 | 19 | 2 | 17 | 595 | 497 | 1075 | 0 | 957 | 97 |

**Supplemental Table S4 (continued).** Antarctic bacteria from 20-ft (TT-abbreviated) and 60-ft BLASTn results (9/17).

| Isolate | Result_ID | Result_Sci_Names | Subject_Blast_Names | %ID | ALIGN_LEN | Mismatches | Gap Opens | Q. start | Q. end | S. start | S. end | Evalue | Bit Score | % Query Coverage Per Subject |
|---|---|---|---|---|---|---|---|---|---|---|---|---|---|---|
| TT-119 | gi\|265678925\|ref\|NR_029233.1\| | *Sporosarcina globispora* | firmicutes | 98.065 | 982 | 15 | 4 | 15 | 992 | 551 | 1532 | 0 | 1677 | 89 |
| TT-120 | gi\|636558971\|ref\|NR_115028.1\| | *Agrococcus jenensis* | high G+C Gram-positive bacteria | 98.125 | 960 | 16 | 2 | 15 | 972 | 514 | 1473 | 0 | 1652 | 87 |
| TT-121 | gi\|631251707\|ref\|NR_112905.1\| | *Tomitella biformata* AHU 1821 | high G+C Gram-positive bacteria | 96.872 | 959 | 25 | 4 | 16 | 970 | 510 | 1467 | 0 | 1590 | 86 |
| TT-137 | gi\|636560215\|ref\|NR_116275.1\| | *Rhodococcus cercidiphylli* | high G+C Gram-positive bacteria | 95.345 | 580 | 22 | 2 | 18 | 596 | 499 | 1074 | 0 | 935 | 97 |
| TT-140 | gi\|636558971\|ref\|NR_115028.1\| | *Agrococcus jenensis* | high G+C Gram-positive bacteria | 97.808 | 958 | 17 | 4 | 19 | 973 | 517 | 1473 | 0 | 1629 | 89 |
| TT-141 | gi\|631252554\|ref\|NR_113752.1\| | *Sporosarcina psychrophila* | firmicutes | 98.283 | 932 | 11 | 3 | 20 | 947 | 545 | 1475 | 0 | 1607 | 89 |
| TT-169 | gi\|636560215\|ref\|NR_116275.1\| | *Rhodococcus cercidiphylli* | high G+C Gram-positive bacteria | 97.064 | 579 | 13 | 3 | 19 | 596 | 498 | 1073 | 0 | 964 | 97 |
| TT-170 | gi\|636560215\|ref\|NR_116275.1\| | *Rhodococcus cercidiphylli* | high G+C Gram-positive bacteria | 97.237 | 579 | 14 | 2 | 18 | 595 | 498 | 1075 | 0 | 973 | 97 |
| TT-171 | gi\|343202628\|ref\|NR_043009.1\| | *Rhodococcus yunnanensis* | high G+C Gram-positive bacteria | 96.366 | 963 | 25 | 8 | 17 | 973 | 507 | 1465 | 0 | 1569 | 87 |

**Supplemental Table S4 (continued).** Antarctic bacteria from 20-ft (TT-abbreviated) and 60-ft BLASTn results (10/17).

| Isolate | Result_ID | Result_Sci_Names | Subject_Blast_Names | %ID | ALIGN_LEN | Mismatches | Gap Opens | Q. start | Q. end | S. start | S. end | Evalue | Bit Score | % Query Coverage Per Subject |
|---|---|---|---|---|---|---|---|---|---|---|---|---|---|---|
| TT-172 | gi\|636560215\|ref\|NR_116275.1\| | *Rhodococcus cercidiphylli* | high G+C Gram-positive bacteria | 96.103 | 975 | 31 | 7 | 18 | 989 | 498 | 1468 | 0 | 1584 | 87 |
| TT-173 | gi\|631251707\|ref\|NR_112905.1\| | *Tomitella biformata* AHU 1821 | high G+C Gram-positive bacteria | 96.367 | 578 | 18 | 3 | 18 | 594 | 510 | 1085 | 0 | 944 | 97 |
| TT-176 | gi\|631251707\|ref\|NR_112905.1\| | *Tomitella biformata* AHU 1821 | high G+C Gram-positive bacteria | 94.71 | 964 | 43 | 6 | 14 | 970 | 507 | 1469 | 0 | 1514 | 88 |
| TT-177 | gi\|1441204194\|ref\|NR_157609.1\| | *Bacillus haynesii* | firmicutes | 99.036 | 934 | 5 | 3 | 20 | 949 | 567 | 1500 | 0 | 1638 | 89 |
| TT-178 | gi\|1040567072\|ref\|NR_137421.1\| | *Bacillus paralicheniformis* | firmicutes | 98.577 | 984 | 8 | 5 | 17 | 995 | 345 | 1327 | 0 | 1702 | 94 |
| TT-179 | gi\|636560215\|ref\|NR_116275.1\| | *Rhodococcus cercidiphylli* | high G+C Gram-positive bacteria | 96.379 | 580 | 18 | 2 | 18 | 596 | 498 | 1075 | 0 | 957 | 97 |
| TT-180 | gi\|636560215\|ref\|NR_116275.1\| | *Rhodococcus cercidiphylli* | high G+C Gram-positive bacteria | 98.826 | 511 | 5 | 1 | 19 | 528 | 498 | 1008 | 0 | 893 | 95 |
| TT-183 | gi\|1779814847\|ref\|NR_164922.1\| | *Ornithinimicrobium cavernae* | high G+C Gram-positive bacteria | 93.473 | 766 | 47 | 2 | 27 | 791 | 573 | 1336 | 0 | 1176 | 73 |
| 202 | gi\|645319993\|ref\|NR_117306.1\| | *Dietzia alimentaria* 72 | high G+C Gram-positive bacteria | 99.15 | 941 | 6 | 2 | 22 | 960 | 485 | 1425 | 0 | 1657 | 84 |
| 209 | gi\|559795330\|ref\|NR_104923.1\| | *Sporosarcina pasteurii* | firmicutes | 95.714 | 980 | 30 | 10 | 22 | 996 | 504 | 1476 | 0 | 1561 | 88 |

**Supplemental Table S4 (continued).** Antarctic bacteria from 20-ft (TT-abbreviated) and 60-ft BLASTn results (11/17).

| Isolate | Result_ID | Result_Sci_Names | Subject_Blast_Names | %ID | ALIGN_LEN | Mismatches | Gap Opens | Q. start | Q. end | S. start | S. end | Evalue | Bit Score | % Query Coverage Per Subject |
|---|---|---|---|---|---|---|---|---|---|---|---|---|---|---|
| 34 | gi\|343200672\|ref\|NR_041359.1\| | *Sporosarcina saromensis* | firmicutes | 98.75 | 960 | 9 | 3 | 18 | 975 | 543 | 1501 | 0 | 1673 | 98 |
| 62 | gi\|219856870\|ref\|NR_024689.1\| | *Bacillus atrophaeus* | firmicutes | 99.485 | 971 | 3 | 2 | 17 | 985 | 544 | 1514 | 0 | 1724 | 98 |
| 92 | gi\|343200672\|ref\|NR_041359.1\| | *Sporosarcina saromensis* | firmicutes | 97.179 | 957 | 24 | 3 | 18 | 971 | 543 | 1499 | 0 | 1612 | 96 |
| 129 | gi\|219857461\|ref\|NR_025049.1\| | *Sporosarcina aquimarina* | firmicutes | 99.17 | 964 | 7 | 1 | 17 | 979 | 545 | 1508 | 0 | 1706 | 97 |
| 149 | gi\|343201095\|ref\|NR_041782.1\| | *Sporosarcina ureae* | firmicutes | 98.017 | 958 | 15 | 4 | 21 | 974 | 545 | 1502 | 0 | 1638 | 98 |
| 150 | gi\|343200672\|ref\|NR_041359.1\| | *Sporosarcina saromensis* | firmicutes | 98.636 | 953 | 10 | 3 | 16 | 966 | 541 | 1492 | 0 | 1655 | 98 |
| 151 | gi\|631253051\|ref\|NR_114249.1\| | *Sporosarcina saromensis* | firmicutes | 98.503 | 935 | 8 | 5 | 17 | 946 | 543 | 1476 | 0 | 1610 | 98 |
| 158 | gi\|631252554\|ref\|NR_113752.1\| | *Sporosarcina psychrophila* | firmicutes | 99.441 | 894 | 4 | 1 | 15 | 907 | 541 | 1434 | 0 | 1587 | 98 |
| 159 | gi\|343201095\|ref\|NR_041782.1\| | *Sporosarcina ureae* | firmicutes | 98.956 | 958 | 7 | 3 | 18 | 972 | 540 | 1497 | 0 | 1679 | 98 |
| 160 | gi\|343201095\|ref\|NR_041782.1\| | *Sporosarcina ureae* | firmicutes | 98.854 | 960 | 7 | 4 | 17 | 972 | 544 | 1503 | 0 | 1674 | 96 |
| 161 | gi\|343200672\|ref\|NR_041359.1\| | *Sporosarcina saromensis* | firmicutes | 98.75 | 960 | 9 | 3 | 20 | 976 | 544 | 1503 | 0 | 1673 | 94 |
| 170 | gi\|343201095\|ref\|NR_041782.1\| | *Sporosarcina ureae* | firmicutes | 99.688 | 963 | 2 | 1 | 16 | 977 | 540 | 1502 | 0 | 1721 | 94 |

**Supplemental Table S4 (continued).** Antarctic bacteria from 20-ft (TT-abbreviated) and 60-ft BLASTn results (12/17).

| Isolate | Result_ID | Result_Sci_Names | Subject_Blast_Names | %ID | ALIGN_LEN | Mismatches | Gap Opens | Q. start | Q. end | S. start | S. end | Evalue | Bit Score | % Query Coverage Per Subject |
|---|---|---|---|---|---|---|---|---|---|---|---|---|---|---|
| 180 | gi\|343201613\|ref\|NR_042339.1\| | *Bacillus aerophilus* | firmicutes | 98.557 | 970 | 8 | 5 | 17 | 980 | 562 | 1531 | 0 | 1675 | 95 |
| 181 | gi\|343200672\|ref\|NR_041359.1\| | *Sporosarcina saromensis* | firmicutes | 98.233 | 962 | 12 | 4 | 17 | 974 | 543 | 1503 | 0 | 1654 | 97 |
| 182 | gi\|343201095\|ref\|NR_041782.1\| | *Sporosarcina ureae* | firmicutes | 98.854 | 960 | 5 | 6 | 15 | 970 | 541 | 1498 | 0 | 1665 | 98 |
| 183 | gi\|343201095\|ref\|NR_041782.1\| | *Sporosarcina ureae* | firmicutes | 99.341 | 910 | 4 | 2 | 17 | 924 | 543 | 1452 | 0 | 1611 | 97 |
| 184 | gi\|343200672\|ref\|NR_041359.1\| | *Sporosarcina saromensis* | firmicutes | 99.168 | 962 | 7 | 1 | 12 | 972 | 541 | 1502 | 0 | 1700 | 98 |
| 185 | gi\|343200672\|ref\|NR_041359.1\| | *Sporosarcina saromensis* | firmicutes | 98.956 | 958 | 7 | 3 | 18 | 972 | 545 | 1502 | 0 | 1676 | 94 |
| 187 | gi\|343201095\|ref\|NR_041782.1\| | *Sporosarcina ureae* | firmicutes | 98.854 | 960 | 8 | 3 | 18 | 974 | 544 | 1503 | 0 | 1679 | 94 |
| 188 | gi\|343200672\|ref\|NR_041359.1\| | *Sporosarcina saromensis* | firmicutes | 97.447 | 940 | 20 | 4 | 17 | 952 | 543 | 1482 | 0 | 1588 | 92 |
| 189 | gi\|343201095\|ref\|NR_041782.1\| | *Sporosarcina ureae* | firmicutes | 98.646 | 960 | 9 | 4 | 18 | 973 | 543 | 1502 | 0 | 1667 | 95 |
| 190 | gi\|559795330\|ref\|NR_104923.1\| | *Sporosarcina pasteurii* | firmicutes | 97.951 | 976 | 16 | 4 | 17 | 989 | 502 | 1476 | 0 | 1662 | 97 |
| 191 | gi\|343201095\|ref\|NR_041782.1\| | *Sporosarcina ureae* | firmicutes | 98.857 | 962 | 8 | 3 | 18 | 977 | 543 | 1503 | 0 | 1682 | 97 |
| 192 | gi\|645319993\|ref\|NR_117306.1\| | *Dietzia alimentaria* 72 | high G+C Gram-positive bacteria | 98.944 | 947 | 7 | 3 | 20 | 965 | 484 | 1428 | 0 | 1658 | 98 |

**Supplemental Table S4 (continued).** Antarctic bacteria from 20-ft (TT-abbreviated) and 60-ft BLASTn results (13/17).

| Isolate | Result_ID | Result_Sci_Names | Subject_Blast_Names | %ID | ALIGN_LEN | Mismatches | Gap Opens | Q. start | Q. end | S. start | S. end | Evalue | Bit Score | % Query Coverage Per Subject |
|---|---|---|---|---|---|---|---|---|---|---|---|---|---|---|
| 193 | gi\|343201095\|ref\|NR_041782.1\| | *Sporosarcina ureae* | firmicutes | 98.859 | 964 | 9 | 2 | 17 | 978 | 544 | 1507 | 0 | 1691 | 98 |
| 195 | gi\|343201095\|ref\|NR_041782.1\| | *Sporosarcina ureae* | firmicutes | 98.749 | 959 | 8 | 4 | 15 | 969 | 544 | 1502 | 0 | 1669 | 94 |
| 197 | gi\|343200672\|ref\|NR_041359.1\| | *Sporosarcina saromensis* | firmicutes | 98.023 | 961 | 16 | 3 | 18 | 975 | 543 | 1503 | 0 | 1648 | 92 |
| 194 | gi\|645319827\|ref\|NR_117184.1\| | *Alkalihalobacillus algicola* | firmicutes | 98.727 | 943 | 7 | 5 | 15 | 955 | 556 | 1495 | 0 | 1635 | 98 |
| 144 | gi\|343201095\|ref\|NR_041782.1\| | *Sporosarcina ureae* | firmicutes | 98.54 | 959 | 12 | 2 | 17 | 973 | 541 | 1499 | 0 | 1665 | 98 |
| 155 | gi\|343202251\|ref\|NR_042537.1\| | *Bhargavaea cecembensis* | firmicutes | 99.033 | 931 | 5 | 4 | 15 | 941 | 559 | 1489 | 0 | 1629 | 98 |
| 196 | gi\|343200588\|ref\|NR_041275.1\| | *Mesobacillus boroniphilus* | firmicutes | 97.79 | 905 | 18 | 2 | 19 | 922 | 565 | 1468 | 0 | 1536 | 98 |
| 199 | gi\|343200672\|ref\|NR_041359.1\| | *Sporosarcina saromensis* | firmicutes | 99.056 | 953 | 7 | 2 | 18 | 969 | 544 | 1495 | 0 | 1675 | 96 |
| 200 | gi\|631251646\|ref\|NR_112844.1\| | *Sporosarcina luteola* | firmicutes | 99.167 | 960 | 5 | 3 | 18 | 974 | 543 | 1502 | 0 | 1690 | 96 |
| 203 | gi\|265678925\|ref\|NR_029233.1\| | *Sporosarcina globispora* | firmicutes | 98.557 | 970 | 13 | 1 | 14 | 982 | 549 | 1518 | 0 | 1687 | 98 |
| 204 | gi\|631251646\|ref\|NR_112844.1\| | *Sporosarcina luteola* | firmicutes | 98.751 | 961 | 9 | 3 | 15 | 972 | 541 | 1501 | 0 | 1676 | 97 |

**Supplemental Table S4 (continued).** Antarctic bacteria from 20-ft (TT-abbreviated) and 60-ft BLASTn results (14/17).

| Isolate | Result_ID | Result_Sci_Names | Subject_Blast_Names | %ID | ALIGN_LEN | Mismatches | Gap Opens | Q. start | Q. end | S. start | S. end | Evalue | Bit Score | % Query Coverage Per Subject |
|---|---|---|---|---|---|---|---|---|---|---|---|---|---|---|
| 205 | gi\|343200672\|ref\|NR_041359.1\| | *Sporosarcina saromensis* | firmicutes | 98.963 | 964 | 7 | 3 | 15 | 977 | 541 | 1502 | 0 | 1686 | 97 |
| 206 | gi\|343201095\|ref\|NR_041782.1\| | *Sporosarcina ureae* | firmicutes | 99.167 | 960 | 6 | 2 | 18 | 976 | 544 | 1502 | 0 | 1694 | 98 |
| 207 | gi\|343200672\|ref\|NR_041359.1\| | *Sporosarcina saromensis* | firmicutes | 98.826 | 937 | 10 | 1 | 18 | 953 | 543 | 1479 | 0 | 1644 | 98 |
| 208 | gi\|343200672\|ref\|NR_041359.1\| | *Sporosarcina saromensis* | firmicutes | 98.938 | 942 | 9 | 1 | 15 | 955 | 541 | 1482 | 0 | 1656 | 95 |
| 210 | gi\|343200672\|ref\|NR_041359.1\| | *Sporosarcina saromensis* | firmicutes | 95.359 | 948 | 41 | 3 | 15 | 960 | 542 | 1488 | 0 | 1521 | 95 |
| 211 | gi\|343200672\|ref\|NR_041359.1\| | *Sporosarcina saromensis* | firmicutes | 98.549 | 965 | 10 | 4 | 16 | 977 | 541 | 1504 | 0 | 1670 | 94 |
| 212 | gi\|343200672\|ref\|NR_041359.1\| | *Sporosarcina saromensis* | firmicutes | 98.853 | 959 | 9 | 2 | 17 | 973 | 543 | 1501 | 0 | 1678 | 94 |
| 214 | gi\|559795330\|ref\|NR_104923.1\| | *Sporosarcina pasteurii* | firmicutes | 97.844 | 974 | 18 | 3 | 16 | 987 | 500 | 1472 | 0 | 1659 | 95 |
| 215 | gi\|343201095\|ref\|NR_041782.1\| | *Sporosarcina ureae* | firmicutes | 98.45 | 968 | 10 | 5 | 17 | 981 | 542 | 1507 | 0 | 1669 | 95 |
| 220 | gi\|265678772\|ref\|NR_029077.1\| | *Alkalihalobacillus algicola* | firmicutes | 97.219 | 971 | 25 | 2 | 28 | 998 | 577 | 1545 | 0 | 1645 | 88 |
| 221 | gi\|343201095\|ref\|NR_041782.1\| | *Sporosarcina ureae* | firmicutes | 97.553 | 940 | 20 | 3 | 18 | 955 | 544 | 1482 | 0 | 1597 | 86 |

**Supplemental Table S4 (continued).** Antarctic bacteria from 20-ft (TT-abbreviated) and 60-ft BLASTn results (15/17).

| Isolate | Result_ID | Result_Sci_Names | Subject_Blast_Names | %ID | ALIGN_LEN | Mismatches | Gap Opens | Q. start | Q. end | S. start | S. end | Evalue | Bit Score | % Query Coverage Per Subject |
|---|---|---|---|---|---|---|---|---|---|---|---|---|---|---|
| 222 | gi\|343201095\|ref\|NR_041782.1\| | *Sporosarcina ureae* | firmicutes | 98.233 | 962 | 12 | 5 | 18 | 976 | 543 | 1502 | 0 | 1652 | 89 |
| 223 | gi\|645321505\|ref\|NR_118455.1\| | *Fictibacillus phosphorivorans* | firmicutes | 99.787 | 469 | 0 | 1 | 17 | 484 | 539 | 1007 | 0 | 838 | 97 |
| 224 | gi\|631252554\|ref\|NR_113752.1\| | *Sporosarcina psychrophila* | firmicutes | 98.387 | 930 | 10 | 4 | 21 | 945 | 544 | 1473 | 0 | 1601 | 90 |
| 227 | gi\|961555219\|ref\|NR_134188.1\| | *Sporosarcina siberiensis* | firmicutes | 90.436 | 941 | 75 | 15 | 17 | 943 | 559 | 1498 | 0 | 1295 | 94 |
| 229 | gi\|343200672\|ref\|NR_041359.1\| | *Sporosarcina saromensis* | firmicutes | 95.228 | 964 | 37 | 9 | 17 | 973 | 543 | 1504 | 0 | 1530 | 87 |
| 232 | gi\|1441204219\|ref\|NR_157634.1\| | *Sporosarcina terrae* | firmicutes | 88.006 | 692 | 76 | 6 | 26 | 714 | 515 | 1202 | 0 | 896 | 71 |
| 233 | gi\|265678925\|ref\|NR_029233.1\| | *Sporosarcina globispora* | firmicutes | 97.248 | 981 | 17 | 7 | 21 | 991 | 553 | 1533 | 0 | 1632 | 89 |
| 234 | gi\|343201095\|ref\|NR_041782.1\| | *Sporosarcina ureae* | firmicutes | 97.597 | 957 | 14 | 8 | 20 | 968 | 543 | 1498 | 0 | 1606 | 89 |
| 236 | gi\|631252554\|ref\|NR_113752.1\| | *Sporosarcina psychrophila* | firmicutes | 98.816 | 929 | 9 | 2 | 21 | 948 | 545 | 1472 | 0 | 1627 | 87 |
| 238 | gi\|219878434\|ref\|NR_025573.1\| | *Paenisporosarcina macmurdoensis* | firmicutes | 98.725 | 941 | 6 | 5 | 17 | 951 | 538 | 1478 | 0 | 1631 | 91 |
| 242 | gi\|961555219\|ref\|NR_134188.1\| | *Sporosarcina siberiensis* | firmicutes | 95.957 | 940 | 32 | 4 | 20 | 953 | 564 | 1503 | 0 | 1521 | 90 |

**Supplemental Table S4 (continued).** Antarctic bacteria from 20-ft (TT-abbreviated) and 60-ft BLASTn results (16/17).

| Isolate | Result_ID | Result_Sci_Names | Subject_Blast_Names | %ID | ALIGN_LEN | Mismatches | Gap Opens | Q. start | Q. end | S. start | S. end | Evalue | Bit Score | % Query Coverage Per Subject |
|---|---|---|---|---|---|---|---|---|---|---|---|---|---|---|
| 243 | gi\|265678925\|ref\|NR_029233.1\| | *Sporosarcina globispora* | firmicutes | 96.758 | 987 | 21 | 7 | 16 | 993 | 549 | 1533 | 0 | 1627 | 89 |
| 244 | gi\|343201095\|ref\|NR_041782.1\| | *Sporosarcina ureae* | firmicutes | 96.258 | 962 | 28 | 8 | 16 | 969 | 542 | 1503 | 0 | 1569 | 90 |
| 247 | gi\|631253051\|ref\|NR_114249.1\| | *Sporosarcina saromensis* | firmicutes | 96.739 | 920 | 25 | 5 | 19 | 934 | 542 | 1460 | 0 | 1526 | 86 |
| 248 | gi\|631252554\|ref\|NR_113752.1\| | *Sporosarcina psychrophila* | firmicutes | 97.959 | 931 | 14 | 4 | 18 | 944 | 544 | 1473 | 0 | 1589 | 87 |
| 249 | gi\|343200672\|ref\|NR_041359.1\| | *Sporosarcina saromensis* | firmicutes | 98.644 | 959 | 10 | 3 | 19 | 974 | 545 | 1503 | 0 | 1666 | 89 |
| 251 | gi\|566084948\|ref\|NR_108491.1\| | *Cytobacillus gottheilii* | firmicutes | 97.907 | 908 | 15 | 4 | 16 | 920 | 563 | 1469 | 0 | 1551 | 86 |
| 252 | gi\|631252554\|ref\|NR_113752.1\| | *Sporosarcina psychrophila* | firmicutes | 97.964 | 933 | 15 | 3 | 18 | 946 | 543 | 1475 | 0 | 1596 | 89 |
| 253 | gi\|265678925\|ref\|NR_029233.1\| | *Sporosarcina globispora* | firmicutes | 97.053 | 984 | 23 | 5 | 16 | 994 | 551 | 1533 | 0 | 1639 | 94 |
| 254 | gi\|343202591\|ref\|NR_042967.1\| | *Virgibacillus halodenitrificans* | firmicutes | 93.439 | 945 | 52 | 6 | 26 | 961 | 570 | 1513 | 0 | 1443 | 89 |
| 255 | gi\|631252639\|ref\|NR_113837.1\| | *Sporosarcina globispora* | firmicutes | 79.16 | 619 | 117 | 12 | 56 | 664 | 579 | 1195 | 3.37E-154 | 545 | 91 |

**Supplemental Table S4 (continued).** Antarctic bacteria from 20-ft (TT-abbreviated) and 60-ft BLASTn results (17/17).

| Isolate | Result_ID | Result_Sci_Names | Subject_Blast_Names | %ID | ALIGN_LEN | Mismatches | Gap Opens | Q. start | Q. end | S. start | S. end | Evalue | Bit Score | % Query Coverage Per Subject |
|---|---|---|---|---|---|---|---|---|---|---|---|---|---|---|
| 257 | gi\|631251707\|ref\|NR_112905.1\| | *Tomitella biformata* AHU 1821 | high G+C Gram-positive bacteria | 92.239 | 786 | 55 | 6 | 17 | 800 | 510 | 1291 | 0 | 1159 | 90 |

**Supplemental Table S5.** Gulf of Mexico multinomial logistic regression results.

| **Genera** | **Intercept** | **Media:** High Nutrient, Artificial Seawater | **Method:** Heat-treat | **Temperature:** 4ºC |
|---|---|---|---|---|
| *Alcanivorax* | -2.2533 (*) | -26.7989 (***) | -24.408 (***) | 26.2927 (NA) |
| *Aliihoeflea* | -35.3683 (***) | 15.7359 (NA) | -12.1619 (***) | 42.2254 (***) |
| *Arthrobacter* | -27.0951 (NA) | -1.4466 (NA) | -24.9645 (***) | 51.1345 (**) |
| *Cellulomonas* | -24.7333 (***) | 22.5954 (***) | -14.5577 (***) | -7.1651 (***) |
| *Dietzia* | -2.9616 (*) | 0.1453 (NA) | -22.6715 (***) | 26.3226 (NA) |
| *Enterobacter* | -35.1643 (***) | -14.8024 (***) | 16.8914 (NA) | 41.0044 (***) |
| *Fictibacillus* | -3.5609 (**) | 1.3518 (NA) | 1.3099 (NA) | 25.7717 (NA) |
| *Kocuria* | -2.2532 (*) | -16.2951 (***) | -13.8926 (NA) | -4.4317 (***) |
| *Kytococcus* | -26.2269 (***) | -17.3576 (***) | 24.9792 (***) | -7.9015 (***) |
| *Mesorhizobium* | -0.6793 (NA) | -0.9823 (NA) | -0.4419 (NA) | 24.2142 (NA) |
| *Metabacillus* | -1.9371 (*) | -0.6654 (NA) | -25.1337 (NA) | 25.5114 (NA) |
| *Micrococcus* | -23.876 (NA) | -21.4805 (***) | -22.1704 (***) | 47.9151 (**) |
| *Mycobacterium* | -26.2269 (***) | -17.3576 (***) | 24.9792 (***) | -7.9015 (***) |
| *Mycolicibacterium* | -0.184 (NA) | -1.2207 (NA) | 0.0617 (NA) | -39.5081 (***) |
| *Nocardia* | -91.5069 (NA) | 40.1065 (NA) | 34.0576 (NA) | 58.507 (NA) |
| *Paenibacillus* | -24.7333 (***) | 22.5954 (***) | -14.5577 (***) | -7.1651 (***) |
| *Paenisporosarcina* | -27.0951 (NA) | -1.4466 (NA) | -24.9645 (***) | 51.1344 (**) |
| *Priestia* | -0.6697 (NA) | 0.4959 (NA) | -0.4571 (NA) | -15.2353 (***) |
| *Pseudoalteromonas* | -27.624 (***) | 26.1792 (***) | -21.3221 (***) | -14.7072 (***) |
| *Pseudomonas* | -26.5708 (NA) | -32.5317 (***) | -29.7254 (***) | 51.3032 (**) |
| *Pseudorhizobium* | -3.3502 (*) | 0.8059 (NA) | -29.8997 (***) | 26.983 (NA) |
| *Rhodococcus* | -2.5916 (*) | 0.0223 (NA) | 1.1275 (NA) | 24.5048 (NA) |
| *Rossellomorea* | -0.8669 (NA) | -33.199 (***) | -27.736 (***) | -15.8862 (***) |
| *Streptomyces* | -2.6302 (*) | -0.6659 (NA) | -21.306 (***) | 25.5118 (NA) |
| **Baseline / Reference** | compared to *Bacillus* | compared to Low Nutrient, Artificial Seawater | compared to Stamp | compared to 25ºC |

*Two-tailed z-test p-values: *** p ≤ 0.001, ** p ≤ 0.01, * p ≤ 0.05, Not Available (NA)*

**Supplemental Table S6.** Antarctica multinomial logistic regression results.

| **Genera** | **Intercept** | **Depth:** 20ft | **Media:** High Nutrient, Natural Seawater | **Media:** Low Nutrient, Artificial Seawater | **Method:** Light | **Temperature:** 30ºC | **Temperature:** 4ºC |
|---|---|---|---|---|---|---|---|
| *Agrococcus* | -328.5037 (***) | 324.9277 (***) | 2.0763 (NA) | -110.0627 (***) | 2.279 (NA) | -0.0637 (NA) | -89.8563 (***) |
| *Alkalihalobacillus* | -184.2873 (***) | -68.7374 (***) | 159.812 (***) | -3.0582 (***) | 24.5323 (***) | -152.4036 (NA) | -223.9741 (***) |
| *Bacillus* | -2.0364 (*) | 2.0747 (NA) | -79.4015 (***) | -0.8457 (NA) | -1.5344 (NA) | -110.8949 (NA) | -108.185 (***) |
| *Bhargavaea* | -279.6971 (***) | -88.3586 (***) | -168.642 (***) | -116.3826 (***) | 131.8499 (***) | 147.9381 (***) | -36.6279 (***) |
| *Cytobacillus* | -261.8102 (***) | -31.0107 (***) | 57.3341 (***) | -10.227 (NA) | 121.3829 (***) | 81.3829 (***) | -78.2182 (NA) |
| *Dietzia* | -3.4844 (***) | 2.4245 (**) | -0.1687 (NA) | 0.2262 (NA) | 0.4656 (NA) | 0.1071 (NA) | -194.5197 (NA) |
| *Fictibacillus* | -244.8763 (***) | -38.1318 (NA) | -13.4899 (***) | 113.8469 (***) | -52.0537 (***) | 129.999 (***) | -45.3761 (NA) |
| *Mesobacillus* | -279.6972 (***) | -88.3586 (***) | -168.642 (***) | -116.3826 (***) | 131.8499 (***) | 147.938 (***) | -36.6279 (***) |
| *Metabacillus* | -296.3174 (***) | 147.9795 (***) | -97.8172 (***) | 0.5921 (NA) | -0.1601 (NA) | 147.0385 (***) | -17.2014 (***) |
| *Micrococcus* | -356.5389 (***) | 104.2439 (***) | -101.2743 (***) | -114.1681 (***) | 143.643 (***) | 108.7945 (***) | -16.0997 (***) |
| *Nocardioides* | -215.6112 (***) | 215.5761 (***) | -88.1339 (***) | -146.995 (***) | -81.8681 (***) | -123.8068 (***) | -31.7294 (***) |
| *Ornithinicoccus* | -199.8762 (***) | 199.1754 (***) | 0.5112 (NA) | -120.9515 (***) | -0.5602 (NA) | -143.0063 (***) | -63.474 (***) |
| *Ornithinimicrobium* | -215.6112 (***) | 215.5761 (***) | -88.1339 (***) | -146.995 (***) | -81.8681 (***) | -123.8068 (***) | -31.7294 (***) |
| *Paenisporosarcina* | -221.447 (**) | 87.4228 (NA) | -14.1307 (***) | 84.2778 (NA) | -46.3108 (***) | 37.0994 (NA) | 136.476 (NA) |
| *Psychrobacter* | -235.6525 (***) | 235.4874 (***) | 1.0673 (NA) | -143.0858 (***) | -1.8146 (NA) | -0.8618 (NA) | -75.3969 (***) |
| *Rhodococcus* | -290.4609 (***) | 292.0175 (***) | 0.5981 (NA) | -1.6129 (NA) | -0.9038 (NA) | -2.8507 (**) | -93.7595 (***) |
| *Robertmurraya* | -437.3771 (***) | 104.1499 (***) | -19.1048 (***) | 115.3847 (***) | 95.294 (***) | 122.1967 (***) | -26.7847 (***) |
| *Rossellomorea* | -373.3036 (***) | 112.1518 (***) | -108.2136 (***) | -121.7074 (***) | 148.5075 (***) | 113.1924 (***) | -17.2241 (***) |
| *Tomitella* | -4.3733 (***) | 2.7056 (*) | -0.0454 (NA) | -0.0154 (NA) | 0.3665 (NA) | 0.8079 (NA) | -95.606 (***) |
| *Virgibacillus* | -164.9438 (***) | -77.4883 (***) | -131.2587 (***) | -124.2121 (***) | -50.3707 (***) | 164.3112 (***) | -27.1565 (***) |
| **Baseline / Reference** | compared to *Sporosarcina* | compared to 60ft | compared to High Nutrient, Artificial Seawater | | compared to Dark | compared to 25ºC | |

*Two-tailed z-test p-values: *** p ≤ 0.001, ** p ≤ 0.01, * p ≤ 0.05, Not Available (NA)*
